# Repurposing antihypertensive, lipid-lowering and antidiabetic drugs for lacunar stroke

**DOI:** 10.1101/2022.11.28.518274

**Authors:** Linjing Zhang, Kailin Xia, Zhou Yu, Yu Fu, Tao Huang, Dongsheng Fan

**Author notes:** **Corresponding author (T.H. and D.F. contributed equally):** Dongsheng Fan: Department of Neurology, Peking University Third Hospital, Beijing, China, Tao Huang, Department of Epidemiology and Biostatistics, School of Public Health, Peking University, Beijing, China.

## Abstract

**Background:** To estimate the causal associations of modifiable risk factors with lacunar stroke (LS) and repurposing of common antihypertensive, lipid-lowering and antidiabetic drugs to prevent LS.

**Methods:** The effects of common antihypertensive, lipid-lowering and antidiabetic drugs on LS were estimated using a drug-target Mendelian randomization (MR) approach. LS data for the transethnic analysis were derived from meta-analyses comprising 7,338 cases and 254,798 controls.

**Findings:** Genetically predicted hypertension and type 2 diabetes significantly increased LS risk. Elevated triglyceride and apolipoprotein B levels caused a 14% increased LS risk, while elevated apolipoprotein A-I and high-density lipoprotein levels caused a 12% decreased risk. Elevated triglyceride levels remained significantly associated with a higher LS risk in multivariable MR analysis (OR, 1.21; 95% CI, 1.06-1.40, P =0.005). Drug-target MR demonstrated that genetic variants mimicking calcium channel blockers most stably prevented LS (OR, 0.75; 95% CI, 0.61-0.92, P =0.006). The genetic variants at or near *HMGCR* (i.e., mimicking the effect of statins), NPC1L1 (mimicking the effects of ezetimibe) and APOC3 (mimicking antisense anti-apoC3 agents) were predicted to decrease LS incidence.

Genetically proxied GLP1R agonism showed a marginal effect on LS, while a genetically proxied improvement in overall glycemic control was associated with a reduced LS risk (OR, 0.94; 95% CI, 0.92–0.96; *P*=4.58×10^−7^).

**Interpretation:** Repurposing several drugs with well-established safety and low costs for LS prevention in clinical practice may contribute to healthier brain aging.

**Funding:** This study was supported by National Natural Science Foundation of China (grant numbers 8210051863).

## Introduction

Lacunar stroke (LS) is a small subcortical infarct that arises from ischemia in the territory of the deep perforating arteries of the brain ^1^. LS accounts for one-quarter of the overall number of ischemic strokes with long-term intellectual and physical disabilities. However, no preventive treatments for LS are available beyond the management of vascular risk factors, such as blood pressure or diabetes, without specific drug classification. Recently, Matthew Traylor et al. made substantial progress in identifying the genetic mechanisms underlying lacunar stroke by genome-wide association studies (GWAS) ^2^, paving the way for repurposing some drugs to more precisely prevent LS via its pathogenesis. Drug repurposing, otherwise known as drug repositioning, is a strategy that seeks to identify new indications and targets for approved drugs that are beyond the scope of their original medical indications ^3^.

The Mendelian randomization (MR) approach is a widely used genetic epidemiological method for assessing causal associations between risk factors and disease by exploiting genetic variants as instrumental variables (IVs) for exposure ^4^. This method leverages the random allocation of genetic variants at conception to reduce any bias due to confounding and reverse causation that can limit causal inference in observational research. MR can be extended to investigate drug effects by leveraging variation in genes (e.g., *HMGCR*) that encode proteins corresponding to drug targets ^5,6^. Specifically, single-nucleotide polymorphisms (SNPs) in or near the *HMGCR* gene were used as proxies for HMGCR inhibition by statins ^7^.

Here, we first exploited a two-sample MR approach to examine the causal associations of modifiable risk factors with LS. Second, multivariable MR was conducted to estimate the direct causal effect of blood pressure and lipids on LS. Third, drug-targeted MR was applied to evaluate several commonly used classes of antihypertensive, lipid-lowering and antidiabetic agents likely to have efficacy in preventing LS. The study design is presented in **Fig. 1**.

**Fig. 1.**
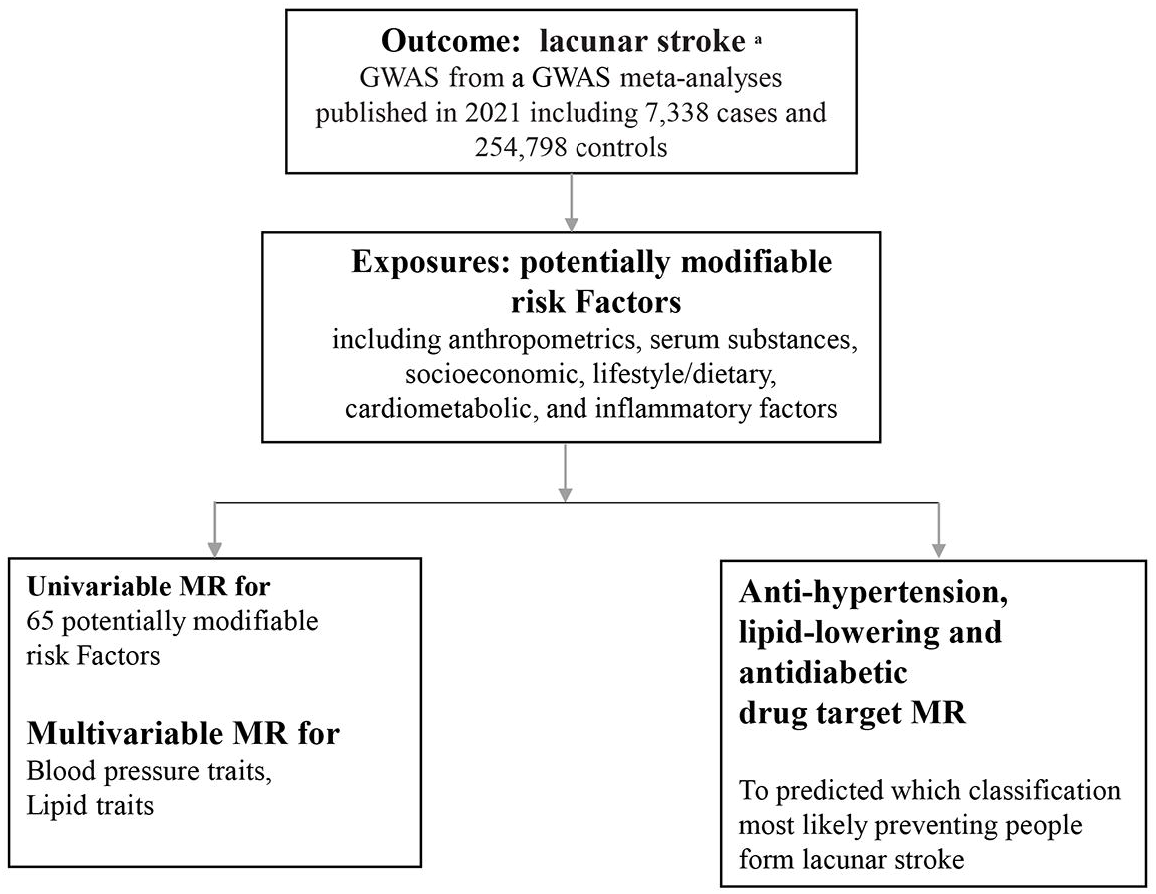
Overall study design. a:Lancet Neurol.2021.May;20(5):351-361. GWAS: genome-wide association study; MR: mendelian randomization.

## MATERIALS AND METHODS

### Modifiable risk factors

We considered potentially modifiable risk factors that can be grouped under the following categories: anthropometry, socioeconomic, lifestyle/dietary, cardiometabolic, etc. These 65 risk factors are listed in **eTable 1**. The eligible risk factors had the most solid evidence from previous observational studies indicating that they may predispose an individual to LS.

### Data sources

We searched PubMed for published genome-wide association studies to obtain the summary data (effect size estimates and their standard errors) of risk factors. We also derived data from MR base https://gwas.mrcieu.ac.uk/. Details on the risk factors that showed significant effects on LS from which we obtained summary data for the current analyses are presented in **eTable 2**.

LS data were derived from meta-analyses from Europe, the USA, and Australia, including previous GWASs and additional cases and controls from the UK DNA LS studies and the

International Stroke Genetics Consortium, comprising a total of 6030 cases and 248□929 controls of European ancestry and 7338 cases and 254□798 controls in the transethnic analysis ^8-11^. These lacunar stroke cases were MRI-confirmed cases, as MRI confirmation of LS is more reliable than standard phenotyping. Genotyping arrays, quality control filters, and imputation reference panels could be found in the study ^2^. Summary-level GWAS data could be derived from https://cd.hugeamp.org/downloads.html.

### Genetic variants

For univariable MR analyses, SNPs with genome-wide significance (P<5×10^−8^) for each risk factor and their effect size estimates and standard errors were also collected. Only independent variants—that is, not in linkage disequilibrium (defined as r^2^<0.001) with other genetic variants for the same risk factor. The IVs (F statistic > 10) for all the exposures were sufficiently informative ^12^. F-statistics were calculated for each variant using the formula F = beta^2^/SE^2^.

For blood pressure (BP)-related (SBP: systolic blood pressure; DBP: diastolic blood pressure; PP: pulse pressure) or lipid-related (ApoB: apolipoprotein B; LDL-C: low-density lipoprotein; TG: triglyceride; ApoA1: apolipoprotein A-I; HDL-C: high-density lipoprotein) traits, one exposure is genetically correlated with other exposures. Thus, we employed multivariable MR (MVMR) analysis to estimate the independent causal effect of each exposure ^13^. As an extension of univariable MR, MVMR concatenating a set of IVs for each exposure estimates the direct effect of exposures on LS risk, whereas univariable MR calculates the total effect.

SNPs for the BP traits were obtained from summary statistics of a large GWAS of BP traits with over 1 million people of European ancestry ^14^. All SNPs with genome-wide significance (n=255, LD_r^2^<0.1) that were associated with any of the BP traits were included in the set of IVs. For the lipid multivariable MR analyses, in Model 1, we pooled all SNPs with genome-wide significance that were associated with any of the traits, including ApoB, LDL-C and TG. In Model 2, the traits included ApoA1 and HDL-C. The IVs were derived from MR base with a threshold clump□r2=0.001 and clump□kb=1000. In all, 384 and 435 IVs were concatenated for Model 1 and Model 2.

For antihypertensive drug-MR, MR analyses were performed to estimate the effect of a 10-mmHg reduction in blood pressure by antihypertensive drugs. Genetic instruments consisted of variants that were associated with each BP trait at genome-wide significance (*P*<5×10^−8^) and located near (+/− 200 kb) or within genes encoding protein targets of 12 antihypertensive medication classes, with effect estimates for each genetic variant derived for each blood pressure trait from the trans-ancestry BP GWAS ^15-17^ (**eTable 3)**. The primary analysis focused on the SBP-lowering effect, with sensitivity analyses considering the remaining BP traits (DBP, PP).

The main method for lipid-lowering drug-MR was to proxy HMG-CoA reductase (mimicking the effect of statins), and 5 SNPs associated with LDL cholesterol at the genome-wide significant level (P□<□5.0□×□10−8) and within ±100 kb windows from the gene region of HMGCR (Entrez Gene: 3156; encoding HMG-CoA reductase) were obtained (**fig. 2**). The variants at or near the *HMGCR, PCSK9*, and *NPC1L1* regions were selected according to the method reported by B. A. Ference ^18^. Similarly, variants at or near the *APOC3* regions were selected using the method proposed by Do et al. ^19^. All variants were chosen based on their associations with the relevant lipid trait (either LDL-C or TG), and they were not strongly correlated (*r*^2^ <0.4 or *r*^2^<0.3) (**eTable 3)**.

**Fig. 2.**
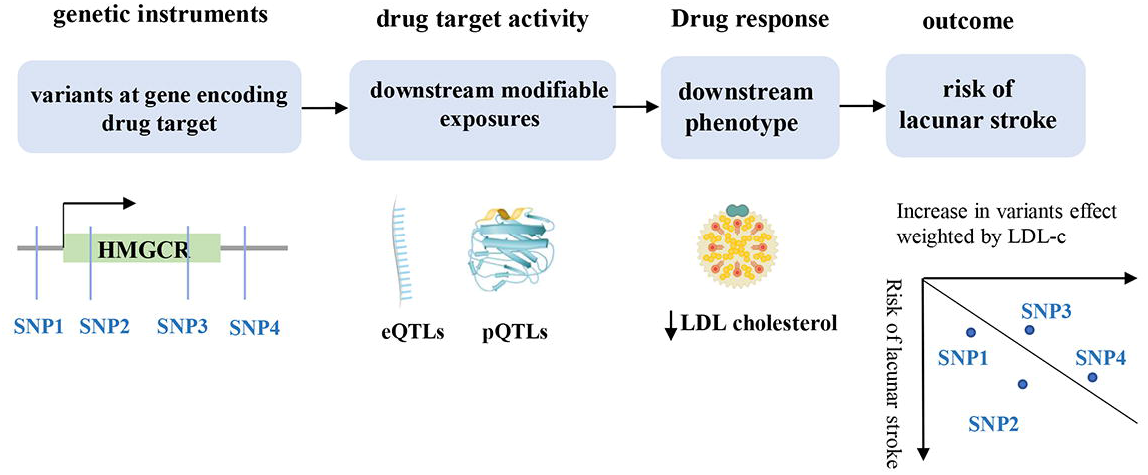
Principles of drug target MR analysis framework. MR: mendelian randomization. SNP: Single Nucleotide Polymorphism; HMGCR: 3-Hydroxy-3-Methylglutaryl-CoA Reductase; eQTLs: expression quantitative trait loci; pQTLs: protein quantitative trait loci; LDL: low density lipoprotein.

Genetic proxies for GLP1R agonism and glycemic control by any mechanism estimated to be associated with glycated hemoglobin (mmol/mol) were derived from 337,000 samples in the UK Biobank ^20,21^. The linkage disequilibrium *r* ^2^ values for variants used as proxies for GLP1R agonism and glycemic control were *r* ^2^<0.1 and 0.001, respectively (**eTable 3)**.

### Mendelian randomization analysis

The principle and main analyses were described in our previous study ^22^. Inverse-variance weighted (IVW) was the primary MR approach in the study. MR□Egger, the weighted median and the simple median were calculated, and the MR□Egger intercept test was used to assess horizontal pleiotropy. We also used the Cochran Q statistic to test for heterogeneity and pleiotropy. For instruments with only 1 variant, Wald-ratio MR was performed.

Next, we conducted multivariable MR and used the multivariable IVW method as the primary approach. To correct for both measured and unmeasured pleiotropy, we also applied multivariable MR□Egger ^23^ and the MR-Lasso method ^24^. We calculated the Cochran Q statistic and multivariable MR□Egger test (intercept) ^13^ to test for heterogeneity and pleiotropy.

The analyses were performed with R version 4.1.1 (R foundation) and the “TwoSample MR” (version 0.5.6) and “MendelianRandomization” (version 0.5.1) packages in R (version 4.1.1). Given that there was only one outcome under investigation (LS), we used a 2-tailed *P* value <.05 to denote evidence against the null hypothesis (i.e., *P* <.05 provided evidence in favor of an association between the exposure and outcome). All human research was approved by the relevant institutional review boards and conducted according to the Declaration of Helsinki. Ethical approval was obtained from relevant Research Ethics Committees and from the review boards of Peking University Third Hospital.

## Results

Among all 65 risk factors (i.e., anthropometric, serum substances, socioeconomic, lifestyle/dietary, cardiometabolic, and inflammatory factors), not surprisingly, genetically predicted hypertension, hyperlipidemia and type 2 diabetes were identified as the predominant high-risk factors for the development of LS (**fig. 3)**. The odds ratio for LS estimated for a 1-SD increase in predisposition to elevated SBP was 1.06 (IVW methods, 95% CI, 1.03-1.08; P =4.64E-7), and the effect was also validated through analysis estimated for DBP and PP. Genetically predicted 1-SD increases in triglyceride and apolipoprotein B levels showed a causal detrimental effect on LS, with odds ratios of 1.14 (95% CI, 1.01-1.29; P =0.027) and 1.15 (95% CI, 1.01-1.31; P =0.030), respectively. The analysis showed a 12% decreased risk of LS with genetically predicted high levels of apolipoprotein A-I and HDL. A genetic predisposition to type 2 diabetes significantly increased the risk of LS (OR, 1.12; 95% CI, 1.06-1.18; P =1.02E-4). In addition, a high level of fasting proinsulin had a detrimental influence on the risk of LS (1.54, 95% CI, 1.10-2.15; P =0.011). There were causal associations between genetically predicted greater height and higher education level and lower odds of LS, and the odds ratios were 0.88 (95% CI, 0.81-0.97; P =0.011) and 0.55 (95% CI, 0.36-0.84; P =0.006), respectively. We also found that a high level of fibrinogen was associated with an increased risk of LS (2.52, 95% CI, 1.16-5.47; P =0.019), and a high risk of atrial fibrillation was associated with a decreased risk of LS (0.93, 95% CI, 0.86-0.98; P =0.010). No other causal evidence of LS risk was found in the analysis. The main results of significant risk factors in univariable MR are presented in **eTable 4**.

**Fig. 3.**
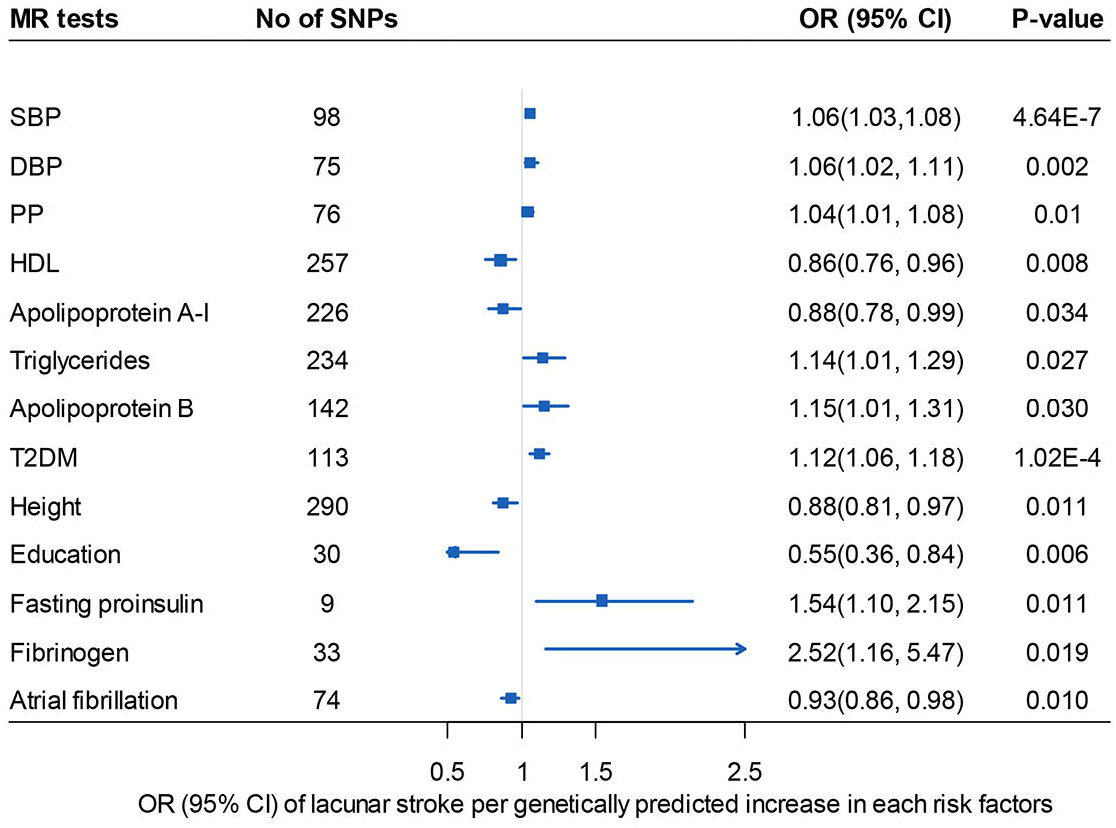
Odds ratios for associations between genetically predicted significant factors and lacunar stroke conduct by univariable MR. MR: mendelian randomization. DBP: diastolic blood pressure; PP: pulse pressure; SBP: systolic blood pressure; HDL: high density lipoprotein; T2DM: diabetes mellitus type 2; OR: odds ratio. 95% CI: 95% confidence interval. HDL, apolipoprotein A-I, triglycerides, apolipoprotein B, height used the multiplicative random effects model due to instrumental heterogeneity (Cochran Q test P< .05).

In blood pressure MVMR analysis, we found little evidence for direct effects of any blood pressure factor on the risk of LS. Specifically, the ORs of LS per 1-SD increase in SBP, DBP and PP were 0.97 (95% CI 0.61–1.56; p = 0.919), 1.08 (95% CI 0.68–1.72; p = 0.729) and 1.07 (95% CI 0.66–1.71; p = 0.789), respectively. For lipid MVMR, when ApoB, LDL-C and TG were assessed together in Model one using the multivariable IVW method, elevated TG levels remained significantly associated with a higher risk of LS (OR, 1.21; 95% CI, 1.06-1.40, P =0.005). In Model two, neither apolipoprotein A-I nor HDL levels showed direct effects on LS risk. The MVMR results are listed in **eTable 5**, and the MR□Egger intercept and Q test results in the study are listed in **eTable 6**.

Furthermore, the antihypertensive drug MR demonstrated that genetic variants mimicking the effect of calcium channel blockers (CCBs) showed the most potent effects in preventing LS. For each 1 SD increase in genetically predicted SBP, the odds ratio of LS was 0.75 (95% CI, 0.61-0.92, P =0.006). The protective effects remained stable in the sensitivity analyses when variants mimicking CCB effects were estimated using DBP and PP (**fig. 4)**. We also found that loop diuretics may prevent LS, as estimated by SBP (OR, 0.50, 95% CI, 0.26-0.97, P =0.041) and DBP (OR, 0.16, 95% CI, 0.04-0.61, P =0.007) (**fig. 4)**. Genetically predicted alpha-adrenoceptor blockers may have marginally protective effects on LS, but only when estimated with SBP (OR, 0.52, 95% CI, 0.27-0.99, P =0.045) (**fig. 4)**. No evidence of efficacy was identified for other antihypertensive drug classes in the analysis.

**Fig. 4.**
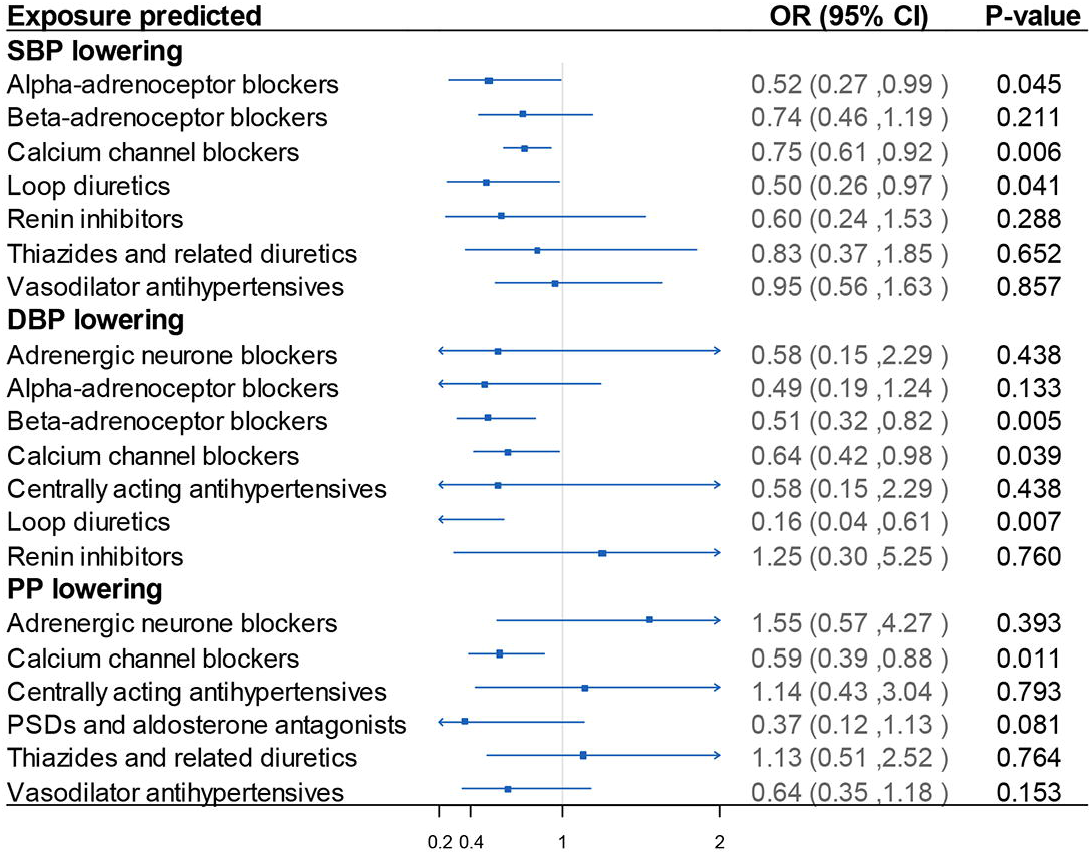
Results of blood pressure-lowering drug target MR analysis. Estimated for SBP lowering target weighted by SBP, DBP lowering target weighted by DBP and PP lowering target weighted by PP. In the left of x-axis=1 presented drug use protective (darked red), in the right of x-axis=1 presented drug use detrimental. MR: mendelian randomization; DBP: diastolic blood pressure; PP: pulse pressure; SBP: systolic blood pressure. OR: odds ratio. 95% CI: 95% confidence interval.

In the LDL-lowering target MR, the LDL-lowering effect predicted by the genetic variants at or near the *HMGCR* gene (i.e., mimicking the effect of statins), NPC1L1 (mimicking the effects of ezetimibe) and APOC3 (mimicking antisense anti-apoC3 agents) may decrease the risk of LS (**fig. 5)**. Genetic proxies for *HMGCR* agonism were associated with a reduced risk of LS (odds ratio per 1 mmol/mol decrease in LDL level 0.53; 95% CI, 0.38-0.73; *P*=8.42×10^−5^), and genetic proxies for *NPC1L1* inhibition were associated with a reduced risk of LS (odds ratio per 1 mmol/mol decrease in LDL level 0.33; 95% CI, 0.18-0.59; *P*=2.36×10^−4^).

**Fig. 5.**
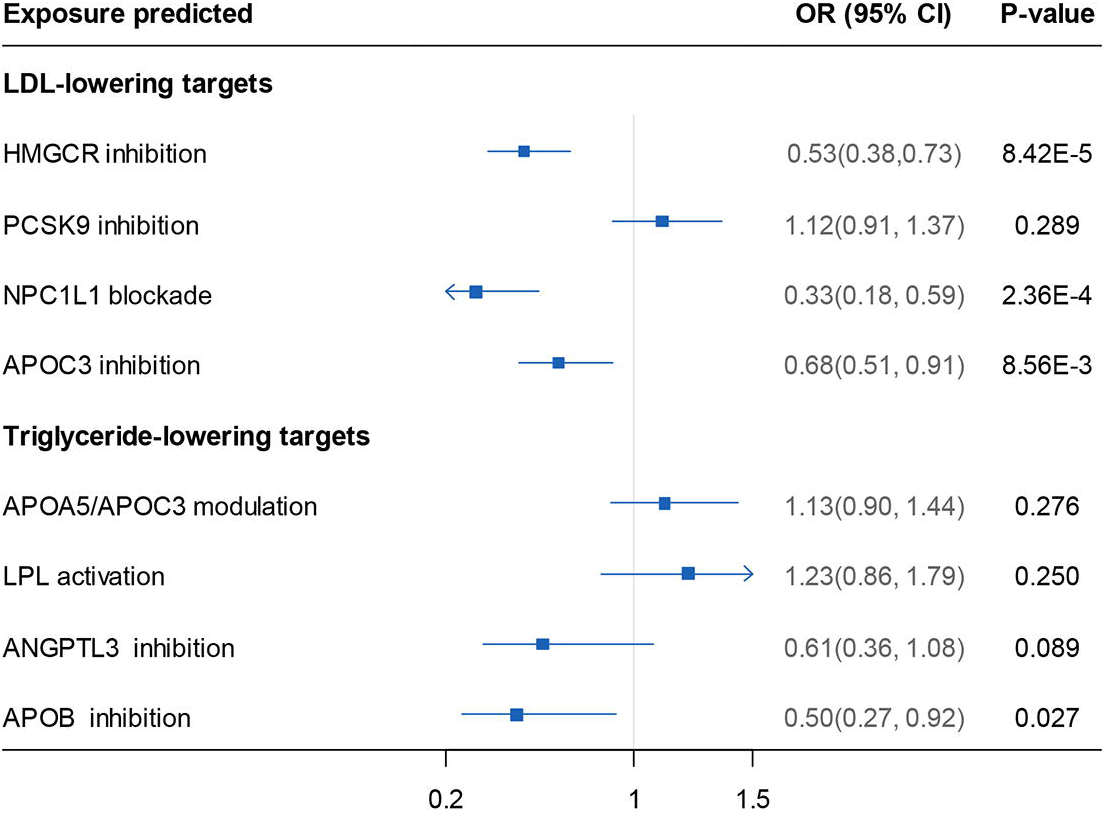
Results of lipid-lowering drug target MR analysis. Estimated for LDL-lowering target weighted by LDL, triglyceride-lowering target weighted by triglyceride. In the left of x-axis=1 presented drug use protective (darked red), in the right of x-axis=1 presented drug use detrimental. MR: mendelian randomization; LDL: low density lipoprotein; HMGCR: 3-Hydroxy-3-Methylglutaryl-CoA Reductase; PCSK9: Proprotein convertase subtilisin kexin type 9; NPC1L1: NPC1 Like Intracellular Cholesterol Transporter 1; APOC3: Apolipoprotein C3; APOA5: Apolipoprotein A5; LPL: Lipoprotein Lipase; ANGPTL3: Angiopoietin Like 3; APOB: Apolipoprotein B; OR: odds ratio. 95% CI: 95% confidence interval.

Significant protective associations of genetically proxied APOC3 inhibition estimated either by LDL or triglyceride lowering with LS risk (0.68, 95% CI, 0.51-0.91; *P*=8.56×10^−3^ and 0.50, 95% CI, 0.27-0.92; *P*=0.027, respectively) were also observed (**fig. 5)**.

We found little evidence for genetic proxies for GLP1R agonism effects on LS; genetically proxied GLP1R agonism showed a marginal effect on LS (OR, 0.47; 95% CI, 0.19-1.12; P =0.088), while a genetically proxied improvement in overall glycemic control was associated with a reduced risk of LS (OR, 0.94; 95% CI, 0.92–0.96; *P*=4.58×10^−7^).

## Discussion

Our MR analysis provided strong genetic evidence that hypertension, hyperlipidemia and type 2 diabetes were the predominant risk factors for the development of LS. Moreover, our MVMR analysis documented that genetically predicted elevated TG levels were still associated with a higher risk of LS. Importantly, the comprehensive drug-target MR approach identified protective effects of common antihypertensive, lipid-lowering medications on LS. CCBs, statins, ezetimibe, and anti-apoC3 agents were most likely to have potential effects of preventing LS. GLP1R agonism did not show such effects, but improvement in overall glycemic control was associated with a reduced risk of LS. This study indicated that old drugs could be repurposed to more precisely prevent LS with substantially lower overall development costs and shorter development timelines ^25^. Such drugs should be given high priority when doctors are making decisions and may contribute to a healthier brain during aging.

Antihypertensive, lipid-lowering agents have been explored with stoke in a previous study. Georgakis MK et al. found that genetic proxies for CCBs showed an inverse association with risk of ischemic stroke compared with proxies for β-blockade, which was particularly strong for small vessel stroke ^26^. Our study showed that among the 12 antihypertensive drug classes, CCBs had the most potent effects in preventing LS and that loop diuretics may prevent the development of LS, as estimated by SBP. β-Blockade did not show protective effects against the risk of LS estimated by SBP. This finding is consistent with results from large-scale meta-analyses of clinical trials and supports that the effects of calcium channel blockers on the risk of stroke are stronger than those for β blockers ^27,28^. Similar analyses have also been performed for lipid-lowering medications. While lipid-lowering variants mimicking statin use have been associated with a lower risk of ischemic stroke, this association was statistically significant only for large artery stroke ^29^. The possible explanation was that the data we used as outcomes made substantial progress in identifying the genetic mechanisms underlying lacunar stroke. We also found little evidence for an effect of PCSK9 inhibitors in preventing LS. Our findings agree with results by Hopewell JC et al. in that PCSK9 inhibitors are unlikely to have an effect on LS risk. Notably, genetic proxies for *HMGCR* agonism, NPC1L1 (mimicking the effects of ezetimibe) and APOC3, were predicted to decrease the risk of LS; they do affect the risk of LS, but they may not do so via LDL. We found little evidence to suggest that LDL itself affects the risk of developing LS. The underlying mechanism needs to be explored in the future.

Blood pressure showed a significant association with LS risk in univariable MR but not in MVMR. A possible explanation for this is that MVMR analysis is used to estimate the independent causal effect of each exposure, whereas univariable MR is used to calculate the total effect. Elevated TG levels were found to be associated with a higher risk of LS conditional on ApoB and LDL-C levels in multivariable MR, which was consistent with the results from univariable MR. This result suggests that compared with ApoB or LDL-C, TG may be more likely to have a detrimental causal effect on LS. Thus, in drug-targeted MR, we applied proxies for lipid-lowering agents estimated by TG-lowering targets.

A strength of our study was the use of two-sample MR, which allows us to utilize the latest GWASs for LS outcomes, which included 7,338 cases and 254,798 controls ^2^. The use of MR over more conventional pharmacoepidemiological approaches also addresses certain forms of confounding. This includes confounding by indication and confounding by the environmental and lifestyle factors of patients, which cannot be fully adjusted for using observational data. This is because measurement error and incomplete capture of all these potential confounding factors inevitably lead to residual confounding. Importantly, drug repurposing, being a less expensive and time-consuming approach, brings effective therapies to patients compared with the cumbersome traditional processes of discovery and development. Moreover, the development risk is significantly lower because repurposing candidates would have already been through several stages of clinical development and have well-established safety and pharmacological profiles, which translates to lower costs and faster development times, thus lowering out-of-pocket costs for patients and ultimately reducing the actual cost of therapy ^30^.

Several limitations merit consideration. We were limited by the fact that MR estimates the effect of lifelong exposure, whereas drugs typically have much shorter periods of exposure, and systolic blood pressure may have age-dependent effects. This means that the effect sizes that we have estimated will not directly reflect what is observed in trials or clinical practice and may not be able to identify critical periods of exposure. Nevertheless, our study assumed a linear relationship between exposures and LS and did not investigate nonlinear effects of the exposures. The populations of exposures and outcomes we explored in the study were not all from subjects of European ancestry, the LS GWAS data were derived from transethnic studies, and the underlying populations were primarily composed of individuals of European ancestry. Thus, bias from population stratification is deemed likely. Finally, completely ruling out pleiotropy or an alternative direct causal pathway is a challenge for all MR analyses because there are probably some unknown confounders that could influence LS. These limitations highlight the importance of cautious interpretation of findings.

## Conclusion

Using an MR design to comprehensively repurpose approved drugs to prevent LS with well-established safety and low costs, this study provided solid evidence for doctors to consider when making decisions in clinical practice and may contribute to a healthier brain during aging.

## Data sharing

All data collected for the study and code used in the analysis will be made available to others.

## Authors and contributors

Linjing Zhang: conceptualisation, data curation, formal analysis, writing – original draft;

Kailin Xia: methodology, software, validation;

Zhou Yu: methodology, software, validation;

Yu Fu: writing– conceptualisation, review & editing;

Tao Huang: conceptualisation, writing– review & editing;

Dongsheng Fan: conceptualisation, writing– review & editing;

