## supplemental table 1-6 for "Repurposing antihypertensive, lipid-lowering and antidiabetic drugs for lacunar stroke"

| **eTable 1. all potential modifaiable risk factors classification groups.** | |
| --- | --- |
| **modifiable risk factors** | **specific exposures** |
| **anthropometry** | waist-to-hip ratio (WHR) |
|  | Body fat |
|  | Height |
|  | Body Mass Index (BMI) |
|  | Bone mineral density |
|  | Childhood BMI |
|  | Birth weight |
| **socioeconomic factors** | Education |
|  | intelligence |
| **lifestyle/dietary factors** | diphenylamine (DPA) |
|  | smoking (number) |
|  | Eicosapntemacnioc Acid (EPA) |
|  | Linoleic acid (LA; 18:2,n6) |
|  | Coffee |
|  | Morning person |
|  | Subjective well-being |
|  | Adrenic acid (22:4,n6) |
|  | Arachidonic acid (AA; 20:4,n6) |
|  | Sedentary |
|  | Carbohydrate |
|  | Gamma linolenic acid (GLA; 18:3,n6) |
| **cadiometabolic factors** | common carotid intima-media thickness |
|  | Coronary heart disease |
|  | HDL cholesterol \|\| id:ieu-b-109 |
|  | Total cholesterol |
|  | homocysteine (Hcy) |
|  | C-reaction protein (CRP) |
|  | Type 2 diabetes |
|  | lipoprotein(a) |
|  | Fasting glucose |
|  | Heart rate |
|  | Fasting proinsulin |
|  | Fasting insulin |
|  | Pulse pressure |
|  | apolipoprotein A-I \|\| id:ieu-b-107 |
|  | diastolic blood pressure (DBP) |
|  | triglycerides \|\| id:ieu-b-111 |
|  | Hypertension |
|  | Fibrinogen |
|  | Adiponectin |
|  | Atrial fibrillation |
|  | Leptin |
|  | LDL cholesterol \|\| id:ieu-b-110 |
|  | HbA1C |
|  | 2h glucose |
|  | systolic blood pressure (SBP) |
|  | apolipoprotein B \|\| id:ieu-b-108 |
| **endogenous substances** | Serum creatinine |
|  | vitamin E |
|  | Uric acid |
|  | eGFRcrea |
|  | Blood urea nitrogen |
|  | Vitamin b12 |
|  | Protein |
|  | vitamin D |
| **neuropsychiatric disorders** | anorexia nervosa |
|  | Schizophrenia |
|  | Neuroticism |
|  | major depression |
|  | Parkinson's disease |
| **other system diseases** | Chronic obstructive pulmonary disease |
|  | Rheumatoid Arthritis |
|  | Osteoporotic fracture |
|  | Crohn's Disease |
|  | Asthma |

| **eTable 2. Summarised GWAS data for potentially modifiable risk factors that shown significant effects on lacunar stroke in univariable MR.** | | | |  |
| --- | --- | --- | --- | --- |
| **Traits** | **No of Ivs in the study (P < 5 × 10-8)** | **Gwas** | **No of sample** | **Units** |
| SBP | 98 | Nat Genet. 2018 Oct; 50(10): 1412–1425. | over one million people of European ancestry | mmHg |
| DBP | 75 |  |  |  |
| PP | 76 |  |  |  |
| HDL cholesterol \|\| id:ieu-b-109 | 257 | UK Biobank, Neale lab, https://gwas.mrcieu.ac.uk/ | 403,943 | mmol/L |
| apolipoprotein A-I \|\| id:ieu-b-107 | 226 |  | 393,193 | mmol/L |
| triglycerides \|\| id:ieu-b-111 | 234 |  | 441,016 | mmol/L |
| apolipoprotein B \|\| id:ieu-b-108 | 142 |  | 439,214 | mmol/L |
| Type 2 diabetes | 113 | Nature Com. 2018;9(1):2941. | 659,316 | increased t2d risk (OR) |
| Height | 290 | Nat Genet. 2014 Nov; 46(11): 1173–1186. | 253,288 | m |
| Education | 30 | Nature. 2016 May 26;533(7604):539-42. | 293,723 | years of schooling |
| Fasting proinsulin | 9 | Diabetes 2011;60(10):2624–2634 | 27,079 | pmol/L |
| Fibrinogen | 33 | Hum Mol Genet. 2016 Jan 15; 25(2): 358–370. | 120,246 | g/l |
| Atrial fibrillation | 90 | Nat Genet. 2018 Sep; 50(9): 1225–1233. | over half a million individuals including 65,446 with AF | increased AF risk (OR) |

| **eTable 3. Genetic variants included in drug-target analyses for each region.** | | | | | | | |
| --- | --- | --- | --- | --- | --- | --- | --- |
| **Traits** | **rsid** | **a1** | **a2** | **a1_freq** | **effect** | **SE** | **p-value** |
| Adrenergic neurone blockers-DBP | rs2692938 | A | G | 0.7879 | -0.0212 | 0.003346 | 2.36E-10 |
| Adrenergic neurone blockers-DBP | rs3111873 | C | G | 0.6675 | 0.02543 | 0.002903 | 1.948E-18 |
| Adrenergic neurone blockers-PP | rs10171471 | C | T | 0.7366 | 0.0283 | 0.003599 | 3.763E-15 |
| Adrenergic neurone blockers-PP | rs3111873 | C | G | 0.6675 | -0.02408 | 0.003342 | 5.803E-13 |
| Adrenergic neurone blockers-PP | rs4526784 | C | G | 0.3742 | 0.02026 | 0.003236 | 3.832E-10 |
| Adrenergic neurone blockers-SBP | rs4526784 | C | G | 0.3742 | 0.02826 | 0.004899 | 8.022E-09 |
| Alpha-adrenoceptor blockers-DBP | rs1741288 | A | G | 0.6384 | -0.01833 | 0.002847 | 1.201E-10 |
| Alpha-adrenoceptor blockers-DBP | rs217728 | T | C | 0.253 | -0.02238 | 0.00314 | 1.018E-12 |
| Alpha-adrenoceptor blockers-DBP | rs2735461 | G | C | 0.9452 | -0.03409 | 0.005994 | 1.291E-08 |
| Alpha-adrenoceptor blockers-DBP | rs415196 | T | C | 0.2749 | 0.01949 | 0.003072 | 2.224E-10 |
| Alpha-adrenoceptor blockers-DBP | rs45471201 | T | C | 0.09837 | -0.02621 | 0.00461 | 1.308E-08 |
| Alpha-adrenoceptor blockers-DBP | rs4872453 | C | T | 0.2532 | -0.01901 | 0.003128 | 1.221E-09 |
| Alpha-adrenoceptor blockers-PP | rs141501760 | T | G | 0.02859 | -0.06011 | 0.009888 | 1.213E-09 |
| Alpha-adrenoceptor blockers-PP | rs217728 | T | C | 0.253 | -0.03471 | 0.003609 | 6.835E-22 |
| Alpha-adrenoceptor blockers-SBP | rs112492 | G | A | 0.2578 | 0.03341 | 0.005404 | 6.329E-10 |
| Alpha-adrenoceptor blockers-SBP | rs141501760 | T | G | 0.02859 | -0.1035 | 0.014939 | 4.271E-12 |
| Alpha-adrenoceptor blockers-SBP | rs217728 | T | C | 0.253 | -0.05864 | 0.005468 | 7.879E-27 |
| Alpha-adrenoceptor blockers-SBP | rs2735461 | G | C | 0.9452 | -0.06705 | 0.010452 | 1.405E-10 |
| Angiotensin converting enzyme inhibitors-DBP | rs4968783 | A | C | 0.6179 | 0.01895 | 0.002808 | 1.494E-11 |
| Angiotensin converting enzyme inhibitors-SBP | rs4968783 | A | C | 0.6179 | 0.02998 | 0.004891 | 8.84E-10 |
| Angiotensin-II receptor antagonists-PP | rs71304101 | A | G | 0.1181 | 0.03005 | 0.004854 | 6.011E-10 |
| Beta-adrenoceptor blockers-DBP | rs11196589 | C | A | 0.3982 | -0.01869 | 0.002815 | 3.127E-11 |
| Beta-adrenoceptor blockers-DBP | rs117624845 | A | G | 0.07281 | -0.04545 | 0.005254 | 5.117E-18 |
| Beta-adrenoceptor blockers-DBP | rs13166730 | T | C | 0.17 | -0.02105 | 0.00375 | 1.984E-08 |
| Beta-adrenoceptor blockers-DBP | rs13242223 | G | A | 0.2339 | -0.01808 | 0.003246 | 2.548E-08 |
| Beta-adrenoceptor blockers-DBP | rs151545 | A | C | 0.1256 | 0.02421 | 0.00428 | 1.545E-08 |
| Beta-adrenoceptor blockers-DBP | rs1741288 | A | G | 0.6384 | -0.01833 | 0.002847 | 1.201E-10 |
| Beta-adrenoceptor blockers-DBP | rs17653278 | G | C | 0.05661 | 0.03425 | 0.005883 | 5.825E-09 |
| Beta-adrenoceptor blockers-DBP | rs180940 | G | A | 0.6693 | -0.02813 | 0.002904 | 3.448E-22 |
| Beta-adrenoceptor blockers-DBP | rs2782980 | C | T | 0.7209 | -0.03911 | 0.003033 | 4.961E-38 |
| Beta-adrenoceptor blockers-DBP | rs2888691 | A | G | 0.1496 | 0.0235 | 0.00384 | 9.346E-10 |
| Beta-adrenoceptor blockers-DBP | rs2907947 | G | A | 0.6218 | 0.0198 | 0.002829 | 2.563E-12 |
| Beta-adrenoceptor blockers-DBP | rs35320559 | C | T | 0.2248 | 0.0211 | 0.003274 | 1.161E-10 |
| Beta-adrenoceptor blockers-DBP | rs3918226 | T | C | 0.07838 | -0.08164 | 0.005151 | 1.49E-56 |
| Beta-adrenoceptor blockers-DBP | rs41313071 | A | C | 0.04352 | 0.04068 | 0.006739 | 1.583E-09 |
| Beta-adrenoceptor blockers-DBP | rs415196 | T | C | 0.2749 | 0.01949 | 0.003072 | 2.224E-10 |
| Beta-adrenoceptor blockers-DBP | rs45471201 | T | C | 0.09837 | -0.02621 | 0.00461 | 1.308E-08 |
| Beta-adrenoceptor blockers-DBP | rs4872453 | C | T | 0.2532 | -0.01901 | 0.003128 | 1.221E-09 |
| Beta-adrenoceptor blockers-DBP | rs68122733 | G | A | 0.1707 | 0.02935 | 0.00363 | 6.195E-16 |
| Beta-adrenoceptor blockers-DBP | rs72829191 | T | C | 0.09773 | -0.02771 | 0.004652 | 2.58E-09 |
| Beta-adrenoceptor blockers-DBP | rs740746 | A | G | 0.7326 | -0.03846 | 0.003084 | 1.092E-35 |
| Beta-adrenoceptor blockers-DBP | rs740956 | C | T | 0.4364 | 0.02149 | 0.002739 | 4.35E-15 |
| Beta-adrenoceptor blockers-DBP | rs741066 | T | C | 0.2942 | -0.01895 | 0.002984 | 2.15E-10 |
| Beta-adrenoceptor blockers-DBP | rs74157560 | T | C | 0.04578 | -0.04864 | 0.006553 | 1.15E-13 |
| Beta-adrenoceptor blockers-DBP | rs75228369 | T | C | 0.04795 | -0.04062 | 0.006536 | 5.152E-10 |
| Beta-adrenoceptor blockers-DBP | rs753482 | A | C | 0.7915 | 0.0249 | 0.003387 | 1.956E-13 |
| Beta-adrenoceptor blockers-DBP | rs77021631 | G | A | 0.1831 | 0.02924 | 0.003519 | 9.571E-17 |
| Beta-adrenoceptor blockers-DBP | rs7737361 | A | G | 0.2028 | 0.02136 | 0.003474 | 7.788E-10 |
| Beta-adrenoceptor blockers-DBP | rs79043825 | A | C | 0.03583 | -0.04211 | 0.007345 | 9.861E-09 |
| Beta-adrenoceptor blockers-DBP | rs855715 | T | G | 0.1177 | 0.04714 | 0.004302 | 6.104E-28 |
| Beta-adrenoceptor blockers-DBP | rs891511 | A | G | 0.3279 | 0.03806 | 0.002942 | 2.771E-38 |
| Beta-adrenoceptor blockers-DBP | rs917875 | C | A | 0.04807 | 0.04424 | 0.006453 | 7.098E-12 |
| Beta-adrenoceptor blockers-PP | rs74157560 | T | C | 0.04578 | -0.04333 | 0.007505 | 7.772E-09 |
| Beta-adrenoceptor blockers-SBP | rs117624845 | A | G | 0.07281 | -0.06348 | 0.009161 | 4.24E-12 |
| Beta-adrenoceptor blockers-SBP | rs13166730 | T | C | 0.17 | -0.04141 | 0.006541 | 2.439E-10 |
| Beta-adrenoceptor blockers-SBP | rs180940 | G | A | 0.6693 | -0.04251 | 0.005057 | 4.244E-17 |
| Beta-adrenoceptor blockers-SBP | rs2782980 | C | T | 0.7209 | -0.05801 | 0.005283 | 4.773E-28 |
| Beta-adrenoceptor blockers-SBP | rs35320559 | C | T | 0.2248 | 0.03729 | 0.005705 | 6.297E-11 |
| Beta-adrenoceptor blockers-SBP | rs3918226 | T | C | 0.07838 | -0.09769 | 0.00899 | 1.679E-27 |
| Beta-adrenoceptor blockers-SBP | rs68122733 | G | A | 0.1707 | 0.04358 | 0.006329 | 5.759E-12 |
| Beta-adrenoceptor blockers-SBP | rs740746 | A | G | 0.7326 | -0.05422 | 0.005371 | 5.809E-24 |
| Beta-adrenoceptor blockers-SBP | rs740956 | C | T | 0.4364 | 0.03184 | 0.004772 | 2.519E-11 |
| Beta-adrenoceptor blockers-SBP | rs741066 | T | C | 0.2942 | -0.02859 | 0.005201 | 3.872E-08 |
| Beta-adrenoceptor blockers-SBP | rs74157560 | T | C | 0.04578 | -0.0936 | 0.011392 | 2.103E-16 |
| Beta-adrenoceptor blockers-SBP | rs753482 | A | C | 0.7915 | 0.03296 | 0.005901 | 2.325E-08 |
| Beta-adrenoceptor blockers-SBP | rs77021631 | G | A | 0.1831 | 0.04808 | 0.006132 | 4.482E-15 |
| Beta-adrenoceptor blockers-SBP | rs855715 | T | G | 0.1177 | 0.06657 | 0.007504 | 7.232E-19 |
| Beta-adrenoceptor blockers-SBP | rs891511 | A | G | 0.3279 | 0.05557 | 0.005123 | 2.061E-27 |
| Calcium channel blockers-DBP | rs10764331 | G | A | 0.418 | -0.02625 | 0.002762 | 2.061E-21 |
| Calcium channel blockers-DBP | rs10828452 | T | A | 0.2049 | 0.01897 | 0.00345 | 3.818E-08 |
| Calcium channel blockers-DBP | rs10828749 | A | G | 0.4104 | 0.02259 | 0.002793 | 6.088E-16 |
| Calcium channel blockers-DBP | rs11062219 | T | C | 0.3678 | -0.02065 | 0.002835 | 3.235E-13 |
| Calcium channel blockers-DBP | rs113210396 | T | G | 0.04499 | 0.03838 | 0.006585 | 5.604E-09 |
| Calcium channel blockers-DBP | rs11591541 | G | A | 0.1571 | -0.02516 | 0.003751 | 1.978E-11 |
| Calcium channel blockers-DBP | rs11720002 | C | T | 0.2723 | -0.01922 | 0.003055 | 3.149E-10 |
| Calcium channel blockers-DBP | rs12258967 | G | C | 0.295 | 0.04153 | 0.002987 | 6.092E-44 |
| Calcium channel blockers-DBP | rs1277754 | G | A | 0.7107 | 0.0182 | 0.003034 | 1.992E-09 |
| Calcium channel blockers-DBP | rs1325990 | G | A | 0.5367 | -0.02532 | 0.00274 | 2.486E-20 |
| Calcium channel blockers-DBP | rs16916944 | T | C | 0.1361 | -0.02668 | 0.004024 | 3.34E-11 |
| Calcium channel blockers-DBP | rs3774751 | T | G | 0.4617 | 0.01918 | 0.002735 | 2.351E-12 |
| Calcium channel blockers-DBP | rs3819531 | T | C | 0.7033 | -0.01745 | 0.002971 | 4.29E-09 |
| Calcium channel blockers-DBP | rs3821843 | A | G | 0.6796 | -0.0255 | 0.002966 | 8.189E-18 |
| Calcium channel blockers-DBP | rs4748478 | A | G | 0.3762 | -0.01853 | 0.002807 | 4.098E-11 |
| Calcium channel blockers-DBP | rs61278674 | G | A | 0.0988 | -0.02709 | 0.004696 | 8.005E-09 |
| Calcium channel blockers-DBP | rs67214975 | A | C | 0.454 | 0.02727 | 0.002742 | 2.666E-23 |
| Calcium channel blockers-DBP | rs7314860 | A | G | 0.1733 | 0.0218 | 0.003654 | 2.434E-09 |
| Calcium channel blockers-DBP | rs7340705 | C | T | 0.3268 | -0.02176 | 0.002931 | 1.13E-13 |
| Calcium channel blockers-DBP | rs76719841 | C | T | 0.0376 | -0.04039 | 0.007201 | 2.036E-08 |
| Calcium channel blockers-PP | rs10764331 | G | A | 0.418 | -0.02422 | 0.003175 | 2.398E-14 |
| Calcium channel blockers-PP | rs10828749 | A | G | 0.4104 | 0.01887 | 0.003212 | 4.215E-09 |
| Calcium channel blockers-PP | rs11248862 | G | A | 0.8731 | 0.02725 | 0.004753 | 9.88E-09 |
| Calcium channel blockers-PP | rs11591541 | G | A | 0.1571 | -0.03572 | 0.004317 | 1.286E-16 |
| Calcium channel blockers-PP | rs12258967 | G | C | 0.295 | 0.03851 | 0.003437 | 3.939E-29 |
| Calcium channel blockers-PP | rs1325990 | G | A | 0.5367 | -0.02449 | 0.003153 | 8.036E-15 |
| Calcium channel blockers-PP | rs13429172 | C | A | 0.3908 | 0.01778 | 0.003224 | 3.484E-08 |
| Calcium channel blockers-PP | rs144399820 | C | T | 0.1404 | -0.02991 | 0.00456 | 5.434E-11 |
| Calcium channel blockers-PP | rs150857355 | C | G | 0.0217 | -0.06581 | 0.010992 | 2.139E-09 |
| Calcium channel blockers-PP | rs1779246 | A | G | 0.8005 | 0.03021 | 0.003956 | 2.226E-14 |
| Calcium channel blockers-PP | rs2497818 | G | A | 0.1066 | -0.03132 | 0.005075 | 6.75E-10 |
| Calcium channel blockers-PP | rs3737984 | T | G | 0.4094 | -0.01829 | 0.003189 | 9.76E-09 |
| Calcium channel blockers-PP | rs3821843 | A | G | 0.6796 | -0.02322 | 0.003414 | 1.044E-11 |
| Calcium channel blockers-PP | rs4748472 | T | C | 0.6621 | -0.01831 | 0.003347 | 4.477E-08 |
| Calcium channel blockers-PP | rs508011 | A | G | 0.3693 | 0.02025 | 0.003274 | 6.224E-10 |
| Calcium channel blockers-PP | rs67214975 | A | C | 0.454 | 0.02678 | 0.003156 | 2.138E-17 |
| Calcium channel blockers-PP | rs6792713 | T | G | 0.01564 | -0.07387 | 0.01241 | 2.635E-09 |
| Calcium channel blockers-PP | rs71384617 | T | C | 0.3557 | 0.01863 | 0.003386 | 3.773E-08 |
| Calcium channel blockers-PP | rs7894090 | A | G | 0.8551 | 0.02856 | 0.004464 | 1.583E-10 |
| Calcium channel blockers-PP | rs7902194 | T | A | 0.7389 | 0.02424 | 0.003615 | 2.023E-11 |
| Calcium channel blockers-PP | rs7920075 | T | C | 0.5289 | 0.018 | 0.003166 | 1.305E-08 |
| Calcium channel blockers-SBP | rs10764331 | G | A | 0.418 | -0.05149 | 0.00481 | 9.835E-27 |
| Calcium channel blockers-SBP | rs10828452 | T | A | 0.2049 | 0.03633 | 0.006013 | 1.52E-09 |
| Calcium channel blockers-SBP | rs10828749 | A | G | 0.4104 | 0.04209 | 0.004865 | 5.097E-18 |
| Calcium channel blockers-SBP | rs11248862 | G | A | 0.8731 | 0.03994 | 0.007188 | 2.749E-08 |
| Calcium channel blockers-SBP | rs113210396 | T | G | 0.04499 | 0.06767 | 0.011487 | 3.84E-09 |
| Calcium channel blockers-SBP | rs11591541 | G | A | 0.1571 | -0.06238 | 0.006535 | 1.363E-21 |
| Calcium channel blockers-SBP | rs116936375 | A | G | 0.03662 | 0.07097 | 0.012751 | 2.606E-08 |
| Calcium channel blockers-SBP | rs12258967 | G | C | 0.295 | 0.08164 | 0.005204 | 1.873E-55 |
| Calcium channel blockers-SBP | rs1277754 | G | A | 0.7107 | 0.03274 | 0.005285 | 5.822E-10 |
| Calcium channel blockers-SBP | rs1325990 | G | A | 0.5367 | -0.05094 | 0.004773 | 1.389E-26 |
| Calcium channel blockers-SBP | rs150857355 | C | G | 0.0217 | -0.1167 | 0.016625 | 2.233E-12 |
| Calcium channel blockers-SBP | rs16916944 | T | C | 0.1361 | -0.04939 | 0.007013 | 1.887E-12 |
| Calcium channel blockers-SBP | rs17123349 | G | A | 0.09325 | -0.04862 | 0.008158 | 2.526E-09 |
| Calcium channel blockers-SBP | rs1779246 | A | G | 0.8005 | 0.04925 | 0.005989 | 1.978E-16 |
| Calcium channel blockers-SBP | rs2497818 | G | A | 0.1066 | -0.05262 | 0.007695 | 8.012E-12 |
| Calcium channel blockers-SBP | rs3819531 | T | C | 0.7033 | -0.02885 | 0.005177 | 2.508E-08 |
| Calcium channel blockers-SBP | rs3821843 | A | G | 0.6796 | -0.0499 | 0.005168 | 4.709E-22 |
| Calcium channel blockers-SBP | rs4748472 | T | C | 0.6621 | -0.0336 | 0.005064 | 3.239E-11 |
| Calcium channel blockers-SBP | rs61278674 | G | A | 0.0988 | -0.0472 | 0.008178 | 7.864E-09 |
| Calcium channel blockers-SBP | rs67214975 | A | C | 0.454 | 0.05505 | 0.004777 | 1.013E-30 |
| Calcium channel blockers-SBP | rs72957281 | C | T | 0.231 | -0.03309 | 0.005625 | 4.05E-09 |
| Calcium channel blockers-SBP | rs7340705 | C | T | 0.3268 | -0.03579 | 0.005107 | 2.406E-12 |
| Calcium channel blockers-SBP | rs76719841 | C | T | 0.0376 | -0.08408 | 0.012556 | 2.141E-11 |
| Calcium channel blockers-SBP | rs7894090 | A | G | 0.8551 | 0.04157 | 0.006758 | 7.691E-10 |
| Calcium channel blockers-SBP | rs7920075 | T | C | 0.5289 | 0.02926 | 0.004793 | 1.027E-09 |
| Centrally acting antihypertensives-DBP | rs2692938 | A | G | 0.7879 | -0.0212 | 0.003346 | 2.36E-10 |
| Centrally acting antihypertensives-DBP | rs3111873 | C | G | 0.6675 | 0.02543 | 0.002903 | 1.948E-18 |
| Centrally acting antihypertensives-PP | rs10171471 | C | T | 0.7366 | 0.0283 | 0.003599 | 3.763E-15 |
| Centrally acting antihypertensives-PP | rs11718509 | A | G | 0.3807 | -0.02143 | 0.003234 | 3.44E-11 |
| Centrally acting antihypertensives-PP | rs3111873 | C | G | 0.6675 | -0.02408 | 0.003342 | 5.803E-13 |
| Loop diuretics-DBP | rs331077 | T | C | 0.425 | -0.02093 | 0.002767 | 3.886E-14 |
| Loop diuretics-DBP | rs7718312 | C | T | 0.2845 | 0.02122 | 0.003018 | 2.059E-12 |
| Loop diuretics-MAP | rs331077 | T | C | 0.425 | -0.02516 | 0.003265 | 1.292E-14 |
| Loop diuretics-MAP | rs7718312 | C | T | 0.2845 | 0.02755 | 0.003561 | 1.033E-14 |
| Loop diuretics-PP | rs2015637 | C | T | 0.1009 | 0.06118 | 0.005184 | 3.868E-32 |
| Loop diuretics-PP | rs35026266 | T | C | 0.2482 | 0.0239 | 0.003628 | 4.504E-11 |
| Loop diuretics-SBP | rs2015637 | C | T | 0.1009 | 0.04711 | 0.007862 | 2.068E-09 |
| Loop diuretics-SBP | rs331077 | T | C | 0.425 | -0.03282 | 0.004822 | 1.003E-11 |
| Loop diuretics-SBP | rs7718312 | C | T | 0.2845 | 0.03923 | 0.00526 | 8.804E-14 |
| PSDs and aldosterone antagonists-PP | rs1058161 | T | C | 0.02403 | 0.06098 | 0.010302 | 3.237E-09 |
| PSDs and aldosterone antagonists-PP | rs2649599 | G | A | 0.7894 | -0.0281 | 0.003948 | 1.096E-12 |
| PSDs and aldosterone antagonists-PP | rs307349 | T | C | 0.9222 | -0.04558 | 0.005923 | 1.42E-14 |
| PSDs and aldosterone antagonists-PP | rs35975487 | G | A | 0.05493 | -0.03947 | 0.006838 | 7.832E-09 |
| PSDs and aldosterone antagonists-SBP | rs307349 | T | C | 0.9222 | -0.05641 | 0.00902 | 4.014E-10 |
| Renin inhibitors-DBP | rs11240656 | A | G | 0.4633 | -0.01534 | 0.002742 | 2.223E-08 |
| Renin inhibitors-DBP | rs16852778 | T | C | 0.1622 | -0.02162 | 0.003713 | 5.785E-09 |
| Renin inhibitors-DBP | rs56305552 | A | G | 0.502 | 0.01499 | 0.002724 | 3.738E-08 |
| Renin inhibitors-SBP | rs11240656 | A | G | 0.4633 | -0.03151 | 0.004776 | 4.188E-11 |
| Renin inhibitors-SBP | rs4293010 | T | G | 0.9235 | -0.0544 | 0.008956 | 1.251E-09 |
| Thiazides and related diuretics-PP | rs12141314 | G | A | 0.1695 | -0.02576 | 0.004166 | 6.262E-10 |
| Thiazides and related diuretics-PP | rs2015637 | C | T | 0.1009 | 0.06118 | 0.005184 | 3.868E-32 |
| Thiazides and related diuretics-PP | rs2474453 | T | G | 0.4938 | 0.02482 | 0.003194 | 7.778E-15 |
| Thiazides and related diuretics-PP | rs35026266 | T | C | 0.2482 | 0.0239 | 0.003628 | 4.504E-11 |
| Thiazides and related diuretics-PP | rs72634852 | T | C | 0.08407 | 0.03435 | 0.005672 | 1.396E-09 |
| Thiazides and related diuretics-SBP | rs12141314 | G | A | 0.1695 | -0.03636 | 0.00631 | 8.311E-09 |
| Thiazides and related diuretics-SBP | rs2015637 | C | T | 0.1009 | 0.04711 | 0.007862 | 2.068E-09 |
| Thiazides and related diuretics-SBP | rs2474453 | T | G | 0.4938 | 0.03381 | 0.004832 | 2.631E-12 |
| Thiazides and related diuretics-SBP | rs3128290 | T | G | 0.6035 | -0.02723 | 0.004857 | 2.07E-08 |
| Vasodilator antihypertensives-DBP | rs41464847 | G | A | 0.4798 | -0.02017 | 0.002723 | 1.3E-13 |
| Vasodilator antihypertensives-PP | rs11024256 | G | T | 0.345 | 0.02403 | 0.00331 | 3.855E-13 |
| Vasodilator antihypertensives-PP | rs214085 | T | C | 0.4113 | -0.02765 | 0.003186 | 4.045E-18 |
| Vasodilator antihypertensives-PP | rs4526784 | C | G | 0.3742 | 0.02026 | 0.003236 | 3.832E-10 |
| Vasodilator antihypertensives-PP | rs56228409 | C | A | 0.1541 | -0.02669 | 0.004435 | 1.756E-09 |
| Vasodilator antihypertensives-PP | rs61755606 | A | G | 0.1079 | 0.03806 | 0.005106 | 9.026E-14 |
| Vasodilator antihypertensives-PP | rs6855875 | T | C | 0.1906 | -0.02854 | 0.004002 | 9.952E-13 |
| Vasodilator antihypertensives-PP | rs7928810 | A | C | 0.6238 | 0.03342 | 0.003248 | 7.92E-25 |
| Vasodilator antihypertensives-SBP | rs11024256 | G | T | 0.345 | 0.03557 | 0.005011 | 1.262E-12 |
| Vasodilator antihypertensives-SBP | rs214085 | T | C | 0.4113 | -0.03797 | 0.004827 | 3.665E-15 |
| Vasodilator antihypertensives-SBP | rs4526784 | C | G | 0.3742 | 0.02826 | 0.004899 | 8.022E-09 |
| Vasodilator antihypertensives-SBP | rs7928810 | A | C | 0.6238 | 0.04449 | 0.004914 | 1.387E-19 |
| **LDL-lowering target weighted by LDL** |  |  |  |  |  |  |  |
| APOC3 | rs10790162 | A | G | 0.1 | 0.2305 | 0.0065 | 1.00E-200 |
| APOC3 | rs603446 | C | T | 0.55 | 0.0502 | 0.0034 | 3.91E-43 |
| NPC1L1 | rs10234070 | T | C | 0.09631 | 0.0295 | 0.0059 | 0.00000152 |
| NPC1L1 | rs217386 | G | A | 0.5923 | 0.0363 | 0.0038 | 1.20E-19 |
| NPC1L1 | rs2300414 | A | G | 0.06992 | 0.0353 | 0.008 | 0.00000545 |
| NPC1L1 | rs7791240 | C | T | 0.09103 | 0.0425 | 0.0065 | 1.84E-10 |
| NPC1L1 | rs2073547 | G | A | 0.195 | 0.0485 | 0.0049 | 1.92E-21 |
| HMGCR | rs12916 | C | T | 0.4314 | 0.0733 | 0.0038 | 7.79E-78 |
| HMGCR | rs17238484 | T | G | 0.2533 | 0.0627 | 0.0062 | 1.35E-21 |
| HMGCR | rs2006760 | G | C | 0.229 | 0.0533 | 0.0076 | 1.67E-13 |
| HMGCR | rs2303152 | A | G | 0.1201 | 0.0423 | 0.0064 | 1.04E-09 |
| HMGCR | rs5909 | A | G | 0.1016 | 0.0617 | 0.0088 | 4.93E-13 |
| HMGCR | rs10066707 | A | G | 0.396 | 0.05 | 0.005 | 3.00E-19 |
| PCSK9 | rs10493176 | T | G | 0.885 | 0.078 | 0.01 | 2.50E-14 |
| PCSK9 | rs11206510 | T | C | 0.846 | 0.083 | 0.005 | 2.38E-53 |
| PCSK9 | rs11206514 | A | C | 0.611 | 0.051 | 0.004 | 1.00E-32 |
| PCSK9 | rs11583974 | A | G | 0.03 | 0.065 | 0.012 | 0.000000004 |
| PCSK9 | rs11591147 | G | T | 0.983 | 0.497 | 0.018 | 8.60E-143 |
| PCSK9 | rs12067569 | A | G | 0.034 | 0.089 | 0.01 | 2.00E-17 |
| PCSK9 | rs2479394 | G | A | 0.285 | 0.039 | 0.004 | 1.60E-19 |
| PCSK9 | rs2479409 | G | A | 0.333 | 0.064 | 0.004 | 2.51E-50 |
| PCSK9 | rs2495477 | T | C | 0.6 | 0.064 | 0.005 | 7.30E-30 |
| PCSK9 | rs572512 | T | C | 0.346 | 0.048 | 0.005 | 5.30E-26 |
| PCSK9 | rs585131 | T | C | 0.815 | 0.064 | 0.005 | 2.70E-35 |
| **triglyceride-lowering target weighted by triglyceride** |  |  |  |  |  |  |  |
| ANGPTL3 | rs4587594 | G | A | 0.69 | 0.069 | 0.004 | 3.50E-82 |
| APOB | rs676210 | G | A | 0.769 | 0.073 | 0.004 | 3.28E-71 |
| APOA5/APOC3 | rs12280753 | T | C | 6.70E-02 | 0.193 | 0.006 | 1.22E-179 |
| APOA5/APOC3 | rs7350481 | T | C | 0.098 | 0.225 | 0.007 | 1.00E-200 |
| LPL | rs12678919 | A | G | 0.879 | 0.17 | 0.006 | 1.82E-199 |
| **antidiabetic drug target** |  |  |  |  |  |  |  |
| GLP1R | rs10305420 | T | C | 0.39 | -0.051 | 0.016 | 1.30x10-3 |
| GLP1R | rs75151020 | C | A | 0.09 | 0.119 | 0.026 | 7.08x10-6 |
| GLP1R | rs2268647 | T | C | 0.52 | 0.066 | 0.015 | 1.51x10-5 |
| **general glycemic control** |  |  |  |  |  |  |  |
| glycemic control | rs7554251 | C | T | 0.73 | 0.036 | 0.017 | 3.39E-02 |
| glycemic control | rs1127215 | T | C | 0.42 | -0.065 | 0.016 | 5.67E-05 |
| glycemic control | rs66464442 | A | C | 0.32 | 0.121 | 0.016 | 2.05E-13 |
| glycemic control | rs1493694 | T | C | 0.11 | 0.146 | 0.025 | 1.05E-08 |
| glycemic control | rs145904381 | C | T | 0.01 | -0.266 | 0.071 | 1.98E-04 |
| glycemic control | rs2297607 | G | A | 0.24 | 0.051 | 0.018 | 4.65E-03 |
| glycemic control | rs6696888 | A | G | 0.68 | -0.048 | 0.016 | 2.75E-03 |
| glycemic control | rs7546252 | G | A | 0.56 | -0.092 | 0.015 | 1.97E-09 |
| glycemic control | rs539515 | C | A | 0.21 | 0.049 | 0.019 | 9.89E-03 |
| glycemic control | rs2816177 | G | A | 0.41 | 0.049 | 0.016 | 2.25E-03 |
| glycemic control | rs41304257 | G | A | 0.28 | -0.042 | 0.017 | 1.34E-02 |
| glycemic control | rs61817176 | C | A | 0.52 | -0.074 | 0.015 | 1.17E-06 |
| glycemic control | rs10916780 | G | A | 0.2 | -0.045 | 0.019 | 1.77E-02 |
| glycemic control | rs340874 | C | T | 0.57 | 0.166 | 0.015 | 2.68E-26 |
| glycemic control | rs1337101 | T | G | 0.32 | -0.095 | 0.016 | 6.05E-09 |
| glycemic control | rs348330 | A | G | 0.63 | -0.119 | 0.016 | 4.90E-13 |
| glycemic control | rs10925635 | C | A | 0.64 | 0.046 | 0.016 | 4.09E-03 |
| glycemic control | rs17261915 | C | T | 0.25 | 0.074 | 0.018 | 4.64E-05 |
| glycemic control | rs3753693 | T | C | 0.41 | -0.066 | 0.016 | 4.38E-05 |
| glycemic control | rs61779284 | A | G | 0.21 | 0.13 | 0.019 | 2.58E-11 |
| glycemic control | rs79090772 | C | T | 0.09 | -0.219 | 0.027 | 3.87E-15 |
| glycemic control | rs2269247 | T | C | 0.18 | -0.056 | 0.02 | 5.15E-03 |
| glycemic control | rs11583755 | C | A | 0.36 | 0.107 | 0.016 | 6.88E-11 |
| glycemic control | rs2613499 | G | A | 0.19 | -0.052 | 0.019 | 6.23E-03 |
| glycemic control | rs10159026 | T | C | 0.25 | -0.062 | 0.018 | 6.08E-04 |
| glycemic control | rs2482506 | G | C | 0.25 | -0.053 | 0.018 | 3.29E-03 |
| glycemic control | rs11196174 | G | A | 0.29 | 0.252 | 0.017 | 4.84E-45 |
| glycemic control | rs149692182 | T | C | 0.02 | 0.309 | 0.053 | 1.11E-08 |
| glycemic control | rs35676242 | A | C | 0.05 | 0.23 | 0.036 | 4.33E-10 |
| glycemic control | rs11257655 | T | C | 0.21 | 0.219 | 0.019 | 2.56E-28 |
| glycemic control | rs946859 | A | G | 0.47 | -0.075 | 0.015 | 8.44E-07 |
| glycemic control | rs3122231 | C | T | 0.65 | 0.05 | 0.016 | 1.83E-03 |
| glycemic control | rs113899647 | T | C | 0.03 | -0.189 | 0.044 | 2.13E-05 |
| glycemic control | rs949693 | A | G | 0.61 | -0.05 | 0.016 | 1.83E-03 |
| glycemic control | rs11592899 | A | G | 0.34 | -0.055 | 0.016 | 6.23E-04 |
| glycemic control | rs2812535 | A | G | 0.62 | 0.069 | 0.016 | 1.98E-05 |
| glycemic control | rs697239 | C | T | 0.46 | -0.105 | 0.015 | 9.28E-12 |
| glycemic control | rs11201992 | A | C | 0.46 | -0.038 | 0.015 | 1.13E-02 |
| glycemic control | rs1111875 | T | C | 0.41 | -0.181 | 0.016 | 2.28E-27 |
| glycemic control | rs66536955 | C | T | 0.26 | 0.044 | 0.017 | 9.63E-03 |
| glycemic control | rs34041345 | G | T | 0.26 | 0.06 | 0.018 | 9.01E-04 |
| glycemic control | rs529623 | C | T | 0.52 | -0.059 | 0.015 | 9.55E-05 |
| glycemic control | rs10893830 | T | C | 0.13 | -0.058 | 0.023 | 1.16E-02 |
| glycemic control | rs10750397 | G | A | 0.72 | -0.04 | 0.017 | 1.85E-02 |
| glycemic control | rs67232546 | T | C | 0.21 | 0.067 | 0.019 | 4.52E-04 |
| glycemic control | rs117316450 | G | C | 0.02 | 0.316 | 0.054 | 9.80E-09 |
| glycemic control | rs757110 | A | C | 0.64 | -0.112 | 0.016 | 9.28E-12 |
| glycemic control | rs11042987 | A | C | 0.58 | -0.034 | 0.016 | 3.33E-02 |
| glycemic control | rs10831668 | T | C | 0.02 | 0.234 | 0.06 | 1.09E-04 |
| glycemic control | rs231362 | G | A | 0.52 | 0.12 | 0.015 | 8.84E-15 |
| glycemic control | rs10767659 | T | G | 0.67 | -0.041 | 0.016 | 1.04E-02 |
| glycemic control | rs60808706 | A | G | 0.05 | -0.227 | 0.035 | 2.40E-10 |
| glycemic control | rs2289488 | C | G | 0.4 | 0.04 | 0.016 | 1.24E-02 |
| glycemic control | rs62618693 | T | C | 0.05 | -0.144 | 0.037 | 1.13E-04 |
| glycemic control | rs523472 | A | G | 0.72 | -0.056 | 0.017 | 1.03E-03 |
| glycemic control | rs7483027 | C | T | 0.38 | -0.061 | 0.016 | 1.54E-04 |
| glycemic control | rs174541 | C | T | 0.36 | -0.098 | 0.016 | 2.07E-09 |
| glycemic control | rs1143756 | G | A | 0.29 | 0.1 | 0.017 | 8.26E-09 |
| glycemic control | rs3918296 | G | C | 0.03 | -0.249 | 0.049 | 5.65E-07 |
| glycemic control | rs11602873 | T | A | 0.16 | -0.187 | 0.021 | 7.97E-18 |
| glycemic control | rs4945090 | A | T | 0.6 | 0.036 | 0.016 | 2.43E-02 |
| glycemic control | rs12802861 | T | C | 0.28 | -0.052 | 0.017 | 2.28E-03 |
| glycemic control | rs10830963 | G | C | 0.28 | 0.297 | 0.017 | 2.61E-61 |
| glycemic control | rs3020069 | A | G | 0.68 | 0.093 | 0.016 | 1.22E-08 |
| glycemic control | rs1426371 | A | G | 0.26 | -0.074 | 0.018 | 4.64E-05 |
| glycemic control | rs79310463 | T | C | 0.13 | 0.104 | 0.023 | 7.91E-06 |
| glycemic control | rs56348580 | C | G | 0.31 | -0.037 | 0.017 | 2.93E-02 |
| glycemic control | rs7975763 | T | C | 0.2 | -0.057 | 0.019 | 2.75E-03 |
| glycemic control | rs11614914 | T | C | 0.33 | 0.078 | 0.016 | 1.54E-06 |
| glycemic control | rs10841886 | C | T | 0.23 | -0.082 | 0.018 | 6.79E-06 |
| glycemic control | rs1480029 | A | G | 0.46 | 0.042 | 0.015 | 5.15E-03 |
| glycemic control | rs3751239 | G | C | 0.2 | -0.16 | 0.019 | 3.68E-16 |
| glycemic control | rs11063018 | C | T | 0.17 | 0.067 | 0.02 | 8.50E-04 |
| glycemic control | rs74862545 | T | C | 0.02 | -0.279 | 0.052 | 1.34E-07 |
| glycemic control | rs2732469 | A | T | 0.43 | -0.258 | 0.015 | 1.57E-59 |
| glycemic control | rs61937817 | G | T | 0.11 | 0.06 | 0.024 | 1.24E-02 |
| glycemic control | rs11173646 | T | A | 0.82 | -0.046 | 0.02 | 2.13E-02 |
| glycemic control | rs2257883 | A | G | 0.13 | 0.15 | 0.023 | 1.93E-10 |
| glycemic control | rs12371967 | C | T | 0.17 | -0.043 | 0.02 | 3.13E-02 |
| glycemic control | rs10879261 | G | T | 0.41 | 0.068 | 0.016 | 2.59E-05 |
| glycemic control | rs11108094 | A | C | 0.07 | 0.099 | 0.03 | 1.01E-03 |
| glycemic control | rs6538805 | C | T | 0.39 | -0.076 | 0.016 | 2.78E-06 |
| glycemic control | rs9587811 | A | C | 0.41 | -0.056 | 0.016 | 4.98E-04 |
| glycemic control | rs314879 | T | C | 0.79 | -0.069 | 0.019 | 3.07E-04 |
| glycemic control | rs34584161 | G | A | 0.24 | -0.063 | 0.018 | 4.98E-04 |
| glycemic control | rs380854 | A | G | 0.58 | -0.058 | 0.016 | 3.14E-04 |
| glycemic control | rs9316500 | G | T | 0.29 | -0.067 | 0.017 | 9.26E-05 |
| glycemic control | rs7991679 | A | T | 0.16 | -0.081 | 0.021 | 1.29E-04 |
| glycemic control | rs1215451 | A | G | 0.29 | -0.131 | 0.017 | 7.45E-14 |
| glycemic control | rs2295388 | A | G | 0.22 | -0.073 | 0.019 | 1.37E-04 |
| glycemic control | rs4906272 | T | C | 0.16 | 0.046 | 0.021 | 2.83E-02 |
| glycemic control | rs12883788 | T | C | 0.46 | 0.06 | 0.015 | 7.31E-05 |
| glycemic control | rs7147483 | C | T | 0.25 | -0.158 | 0.018 | 2.22E-17 |
| glycemic control | rs723355 | A | G | 0.5 | -0.034 | 0.015 | 2.32E-02 |
| glycemic control | rs4902002 | A | G | 0.71 | -0.034 | 0.017 | 4.51E-02 |
| glycemic control | rs242105 | C | A | 0.28 | 0.062 | 0.017 | 2.89E-04 |
| glycemic control | rs7156625 | A | G | 0.22 | 0.037 | 0.019 | 5.11E-02 |
| glycemic control | rs8010382 | G | A | 0.41 | 0.046 | 0.016 | 4.09E-03 |
| glycemic control | rs8043085 | T | G | 0.23 | 0.071 | 0.018 | 9.14E-05 |
| glycemic control | rs11856877 | G | A | 0.11 | 0.071 | 0.024 | 3.15E-03 |
| glycemic control | rs1473781 | A | G | 0.35 | 0.067 | 0.016 | 3.37E-05 |
| glycemic control | rs149336329 | T | G | 0.05 | -0.272 | 0.037 | 8.86E-13 |
| glycemic control | rs7163757 | T | C | 0.43 | -0.042 | 0.015 | 5.15E-03 |
| glycemic control | rs7178762 | T | C | 0.55 | -0.057 | 0.015 | 1.61E-04 |
| glycemic control | rs9479 | G | A | 0.49 | 0.052 | 0.015 | 5.61E-04 |
| glycemic control | rs8033589 | A | G | 0.76 | 0.058 | 0.018 | 1.32E-03 |
| glycemic control | rs12910361 | G | A | 0.71 | 0.161 | 0.017 | 7.02E-20 |
| glycemic control | rs893617 | T | C | 0.72 | -0.136 | 0.017 | 8.84E-15 |
| glycemic control | rs2290202 | T | G | 0.13 | 0.085 | 0.023 | 2.41E-04 |
| glycemic control | rs9927842 | C | T | 0.84 | -0.056 | 0.021 | 7.67E-03 |
| glycemic control | rs8056890 | A | G | 0.36 | 0.105 | 0.016 | 1.50E-10 |
| glycemic control | rs8054556 | A | G | 0.47 | 0.077 | 0.015 | 4.37E-07 |
| glycemic control | rs55857387 | C | T | 0.2 | -0.142 | 0.019 | 3.81E-13 |
| glycemic control | rs8061528 | T | C | 0.21 | 0.092 | 0.019 | 1.80E-06 |
| glycemic control | rs2024449 | C | T | 0.44 | -0.056 | 0.015 | 2.09E-04 |
| glycemic control | rs1421085 | C | T | 0.4 | 0.154 | 0.016 | 1.84E-20 |
| glycemic control | rs56125990 | G | A | 0.15 | 0.065 | 0.021 | 2.02E-03 |
| glycemic control | rs4788815 | T | A | 0.66 | 0.056 | 0.016 | 4.98E-04 |
| glycemic control | rs72802365 | C | G | 0.08 | -0.163 | 0.029 | 3.48E-08 |
| glycemic control | rs2966117 | T | G | 0.48 | 0.059 | 0.015 | 9.55E-05 |
| glycemic control | rs11117364 | G | A | 0.68 | 0.066 | 0.017 | 1.17E-04 |
| glycemic control | rs66461358 | C | T | 0.15 | 0.06 | 0.021 | 4.32E-03 |
| glycemic control | rs12934854 | A | G | 0.17 | 0.043 | 0.02 | 3.13E-02 |
| glycemic control | rs925095 | T | C | 0.39 | -0.075 | 0.016 | 3.72E-06 |
| glycemic control | rs2297508 | G | C | 0.65 | -0.15 | 0.016 | 1.59E-19 |
| glycemic control | rs9913225 | A | G | 0.58 | -0.075 | 0.016 | 3.72E-06 |
| glycemic control | rs1109442 | C | T | 0.47 | 0.071 | 0.015 | 3.01E-06 |
| glycemic control | rs3110641 | G | A | 0.78 | 0.091 | 0.019 | 2.31E-06 |
| glycemic control | rs11651755 | T | C | 0.51 | -0.124 | 0.015 | 1.20E-15 |
| glycemic control | rs3786017 | C | T | 0.11 | 0.054 | 0.025 | 3.05E-02 |
| glycemic control | rs8071043 | C | T | 0.33 | 0.066 | 0.016 | 4.38E-05 |
| glycemic control | rs1905339 | C | T | 0.34 | 0.097 | 0.016 | 2.96E-09 |
| glycemic control | rs35895680 | A | C | 0.33 | -0.089 | 0.016 | 4.76E-08 |
| glycemic control | rs366577 | T | C | 0.6 | -0.051 | 0.016 | 1.49E-03 |
| glycemic control | rs57767539 | A | G | 0.07 | 0.136 | 0.031 | 1.43E-05 |
| glycemic control | rs11658220 | A | G | 0.1 | 0.1 | 0.025 | 7.31E-05 |
| glycemic control | rs12603589 | C | T | 0.19 | 0.103 | 0.02 | 4.02E-07 |
| glycemic control | rs7224711 | T | C | 0.53 | -0.069 | 0.015 | 5.56E-06 |
| glycemic control | rs303760 | T | C | 0.35 | 0.052 | 0.016 | 1.20E-03 |
| glycemic control | rs16965062 | T | C | 0.43 | 0.034 | 0.015 | 2.32E-02 |
| glycemic control | rs7227272 | A | G | 0.1 | -0.062 | 0.026 | 1.70E-02 |
| glycemic control | rs410150 | T | C | 0.8 | -0.047 | 0.019 | 1.33E-02 |
| glycemic control | rs17596995 | A | G | 0.2 | -0.049 | 0.019 | 9.89E-03 |
| glycemic control | rs1517037 | T | C | 0.19 | -0.093 | 0.02 | 4.42E-06 |
| glycemic control | rs6567160 | C | T | 0.23 | 0.097 | 0.018 | 1.19E-07 |
| glycemic control | rs74625348 | C | G | 0.23 | -0.044 | 0.019 | 2.04E-02 |
| glycemic control | rs12963820 | A | T | 0.27 | 0.034 | 0.017 | 4.51E-02 |
| glycemic control | rs7240767 | C | T | 0.39 | 0.063 | 0.016 | 9.39E-05 |
| glycemic control | rs6565922 | T | C | 0.38 | 0.078 | 0.016 | 1.54E-06 |
| glycemic control | rs9384 | T | G | 0.38 | -0.107 | 0.016 | 6.88E-11 |
| glycemic control | rs10404726 | T | C | 0.47 | -0.035 | 0.015 | 1.95E-02 |
| glycemic control | rs58542926 | T | C | 0.08 | 0.139 | 0.029 | 2.27E-06 |
| glycemic control | rs924150 | C | A | 0.39 | -0.087 | 0.016 | 9.23E-08 |
| glycemic control | rs4805881 | C | A | 0.67 | -0.077 | 0.016 | 2.08E-06 |
| glycemic control | rs429358 | C | T | 0.16 | -0.142 | 0.021 | 4.30E-11 |
| glycemic control | rs8107527 | A | G | 0.28 | 0.105 | 0.017 | 1.53E-09 |
| glycemic control | rs9304665 | A | T | 0.77 | 0.103 | 0.018 | 2.01E-08 |
| glycemic control | rs2115107 | A | G | 0.38 | 0.069 | 0.016 | 1.98E-05 |
| glycemic control | rs116843064 | A | G | 0.02 | -0.15 | 0.055 | 6.41E-03 |
| glycemic control | rs34506349 | A | G | 0.04 | -0.099 | 0.038 | 9.17E-03 |
| glycemic control | rs79950062 | C | T | 0.13 | -0.053 | 0.023 | 2.10E-02 |
| glycemic control | rs9308614 | G | A | 0.15 | -0.09 | 0.022 | 5.04E-05 |
| glycemic control | rs6716394 | A | G | 0.54 | -0.045 | 0.015 | 2.75E-03 |
| glycemic control | rs4668483 | G | A | 0.68 | -0.04 | 0.016 | 1.24E-02 |
| glycemic control | rs10184004 | T | C | 0.41 | -0.115 | 0.016 | 2.68E-12 |
| glycemic control | rs13406280 | T | C | 0.49 | -0.047 | 0.015 | 1.78E-03 |
| glycemic control | rs72917531 | A | C | 0.19 | -0.078 | 0.02 | 1.09E-04 |
| glycemic control | rs36051007 | T | C | 0.32 | -0.035 | 0.017 | 3.92E-02 |
| glycemic control | rs67383253 | C | T | 0.37 | -0.035 | 0.016 | 2.85E-02 |
| glycemic control | rs6712905 | C | T | 0.26 | 0.048 | 0.018 | 7.67E-03 |
| glycemic control | rs4482463 | A | C | 0.92 | -0.063 | 0.029 | 2.96E-02 |
| glycemic control | rs34329895 | G | A | 0.6 | -0.063 | 0.016 | 9.39E-05 |
| glycemic control | rs2943650 | T | C | 0.65 | 0.143 | 0.016 | 6.10E-18 |
| glycemic control | rs13415288 | C | T | 0.34 | 0.059 | 0.016 | 2.48E-04 |
| glycemic control | rs34339006 | T | C | 0.39 | 0.092 | 0.016 | 1.72E-08 |
| glycemic control | rs1260326 | C | T | 0.61 | 0.156 | 0.016 | 6.16E-21 |
| glycemic control | rs77165542 | T | C | 0.04 | -0.155 | 0.042 | 2.46E-04 |
| glycemic control | rs921069 | G | A | 0.58 | -0.038 | 0.016 | 1.74E-02 |
| glycemic control | rs76675804 | C | T | 0.1 | -0.311 | 0.026 | 2.67E-30 |
| glycemic control | rs10193538 | T | G | 0.61 | 0.072 | 0.016 | 8.71E-06 |
| glycemic control | rs243018 | G | C | 0.45 | 0.088 | 0.016 | 6.64E-08 |
| glycemic control | rs10188334 | T | C | 0.17 | -0.087 | 0.02 | 1.69E-05 |
| glycemic control | rs12185610 | C | A | 0.41 | -0.063 | 0.016 | 9.39E-05 |
| glycemic control | rs4671799 | G | A | 0.68 | -0.036 | 0.016 | 2.43E-02 |
| glycemic control | rs4832290 | C | T | 0.77 | -0.053 | 0.018 | 3.29E-03 |
| glycemic control | rs6137042 | A | G | 0.2 | -0.05 | 0.019 | 8.50E-03 |
| glycemic control | rs7274134 | T | C | 0.25 | -0.062 | 0.018 | 6.08E-04 |
| glycemic control | rs6059662 | G | A | 0.65 | 0.037 | 0.016 | 2.06E-02 |
| glycemic control | rs2038457 | G | A | 0.81 | 0.041 | 0.02 | 4.00E-02 |
| glycemic control | rs12625671 | C | T | 0.11 | 0.118 | 0.025 | 3.20E-06 |
| glycemic control | rs6066138 | A | G | 0.28 | -0.135 | 0.017 | 1.36E-14 |
| glycemic control | rs6021276 | C | T | 0.64 | -0.074 | 0.016 | 4.96E-06 |
| glycemic control | rs865034 | C | T | 0.66 | 0.04 | 0.016 | 1.24E-02 |
| glycemic control | rs4810145 | C | T | 0.52 | 0.068 | 0.015 | 7.51E-06 |
| glycemic control | rs6011155 | C | T | 0.37 | -0.074 | 0.016 | 4.96E-06 |
| glycemic control | rs2240716 | T | C | 0.3 | 0.074 | 0.017 | 1.66E-05 |
| glycemic control | rs56392746 | A | G | 0.09 | -0.138 | 0.026 | 1.81E-07 |
| glycemic control | rs75307421 | A | G | 0.02 | 0.151 | 0.061 | 1.32E-02 |
| glycemic control | rs138771 | G | A | 0.81 | -0.055 | 0.02 | 5.99E-03 |
| glycemic control | rs1801645 | T | C | 0.74 | -0.059 | 0.018 | 1.09E-03 |
| glycemic control | rs17036126 | T | C | 0.13 | 0.127 | 0.023 | 5.92E-08 |
| glycemic control | rs11708067 | G | A | 0.25 | -0.262 | 0.018 | 1.55E-43 |
| glycemic control | rs17036160 | T | C | 0.12 | -0.088 | 0.024 | 2.69E-04 |
| glycemic control | rs9873519 | T | C | 0.53 | 0.097 | 0.015 | 2.70E-10 |
| glycemic control | rs667920 | T | G | 0.77 | 0.038 | 0.018 | 3.45E-02 |
| glycemic control | rs9289556 | T | C | 0.73 | -0.077 | 0.017 | 7.64E-06 |
| glycemic control | rs56243018 | C | A | 0.05 | -0.214 | 0.036 | 5.82E-09 |
| glycemic control | rs28502438 | C | T | 0.43 | -0.051 | 0.016 | 1.49E-03 |
| glycemic control | rs7633673 | A | G | 0.41 | -0.086 | 0.016 | 1.28E-07 |
| glycemic control | rs11706810 | C | T | 0.48 | -0.106 | 0.015 | 5.99E-12 |
| glycemic control | rs13099581 | T | C | 0.14 | -0.06 | 0.022 | 6.41E-03 |
| glycemic control | rs8192675 | C | T | 0.29 | -0.188 | 0.017 | 2.89E-26 |
| glycemic control | rs6444036 | T | G | 0.16 | 0.041 | 0.021 | 5.05E-02 |
| glycemic control | rs9859406 | A | G | 0.31 | 0.167 | 0.017 | 3.21E-21 |
| glycemic control | rs2041965 | T | C | 0.34 | -0.083 | 0.016 | 3.33E-07 |
| glycemic control | rs6777684 | G | A | 0.61 | 0.134 | 0.016 | 5.25E-16 |
| glycemic control | rs13094957 | C | T | 0.2 | -0.131 | 0.019 | 1.84E-11 |
| glycemic control | rs1470560 | A | G | 0.37 | 0.037 | 0.016 | 2.06E-02 |
| glycemic control | rs2624847 | T | G | 0.74 | -0.084 | 0.017 | 1.12E-06 |
| glycemic control | rs13434089 | C | T | 0.12 | -0.082 | 0.024 | 6.72E-04 |
| glycemic control | rs9870517 | C | A | 0.4 | -0.096 | 0.016 | 4.24E-09 |
| glycemic control | rs1374915 | C | T | 0.42 | -0.036 | 0.016 | 2.43E-02 |
| glycemic control | rs1523766 | G | A | 0.5 | -0.031 | 0.015 | 3.84E-02 |
| glycemic control | rs978444 | T | G | 0.55 | -0.057 | 0.015 | 1.61E-04 |
| glycemic control | rs3872707 | A | G | 0.12 | 0.049 | 0.023 | 3.29E-02 |
| glycemic control | rs7659468 | G | T | 0.49 | -0.103 | 0.015 | 2.20E-11 |
| glycemic control | rs11728350 | G | A | 0.13 | 0.11 | 0.023 | 2.39E-06 |
| glycemic control | rs77141743 | A | G | 0.16 | 0.045 | 0.021 | 3.19E-02 |
| glycemic control | rs730831 | G | T | 0.04 | -0.123 | 0.041 | 2.75E-03 |
| glycemic control | rs2604918 | T | G | 0.33 | -0.063 | 0.016 | 9.39E-05 |
| glycemic control | rs2125799 | C | T | 0.33 | 0.06 | 0.016 | 1.96E-04 |
| glycemic control | rs28819812 | A | C | 0.32 | -0.06 | 0.016 | 1.96E-04 |
| glycemic control | rs4865436 | G | C | 0.29 | 0.05 | 0.018 | 5.51E-03 |
| glycemic control | rs2169033 | T | C | 0.68 | 0.081 | 0.017 | 2.60E-06 |
| glycemic control | rs55691245 | A | G | 0.14 | -0.16 | 0.022 | 1.51E-12 |
| glycemic control | rs7664347 | C | T | 0.64 | -0.04 | 0.016 | 1.24E-02 |
| glycemic control | rs10938398 | A | G | 0.43 | 0.04 | 0.016 | 1.24E-02 |
| glycemic control | rs1996617 | C | T | 0.29 | 0.101 | 0.017 | 5.93E-09 |
| glycemic control | rs114447556 | T | C | 0.08 | 0.08 | 0.029 | 5.84E-03 |
| glycemic control | rs10937721 | C | G | 0.59 | 0.142 | 0.016 | 1.01E-17 |
| glycemic control | rs73222806 | G | C | 0.05 | 0.098 | 0.035 | 5.15E-03 |
| glycemic control | rs6835992 | G | A | 0.69 | 0.066 | 0.017 | 1.17E-04 |
| glycemic control | rs993380 | G | A | 0.67 | -0.059 | 0.016 | 2.48E-04 |
| glycemic control | rs28408270 | T | G | 0.47 | -0.05 | 0.015 | 9.01E-04 |
| glycemic control | rs1961224 | G | A | 0.35 | -0.065 | 0.016 | 5.67E-05 |
| glycemic control | rs141146025 | A | C | 0.02 | 0.129 | 0.058 | 2.59E-02 |
| glycemic control | rs75432112 | A | G | 0.05 | 0.195 | 0.036 | 1.03E-07 |
| glycemic control | rs329118 | T | C | 0.42 | 0.041 | 0.016 | 1.04E-02 |
| glycemic control | rs111686785 | G | A | 0.03 | 0.182 | 0.044 | 4.18E-05 |
| glycemic control | rs72734782 | G | A | 0.21 | 0.066 | 0.019 | 5.47E-04 |
| glycemic control | rs12514030 | G | T | 0.12 | -0.103 | 0.023 | 9.60E-06 |
| glycemic control | rs1650505 | A | G | 0.21 | 0.06 | 0.019 | 1.64E-03 |
| glycemic control | rs4343858 | A | G | 0.23 | -0.042 | 0.018 | 1.95E-02 |
| glycemic control | rs138373837 | T | C | 0.02 | 0.101 | 0.05 | 4.30E-02 |
| glycemic control | rs62366821 | G | A | 0.49 | -0.055 | 0.015 | 2.69E-04 |
| glycemic control | rs10067659 | C | G | 0.79 | -0.081 | 0.019 | 2.45E-05 |
| glycemic control | rs4865796 | A | G | 0.69 | 0.049 | 0.017 | 3.99E-03 |
| glycemic control | rs464605 | T | C | 0.75 | 0.08 | 0.019 | 3.06E-05 |
| glycemic control | rs34341 | T | A | 0.58 | 0.073 | 0.016 | 6.58E-06 |
| glycemic control | rs7732130 | A | G | 0.68 | -0.132 | 0.016 | 1.36E-15 |
| glycemic control | rs6870983 | T | C | 0.21 | -0.067 | 0.019 | 4.52E-04 |
| glycemic control | rs34483452 | A | C | 0.14 | 0.077 | 0.023 | 8.56E-04 |
| glycemic control | rs7752666 | T | C | 0.32 | -0.035 | 0.017 | 3.92E-02 |
| glycemic control | rs80196932 | C | T | 0.16 | -0.064 | 0.021 | 2.36E-03 |
| glycemic control | rs11759026 | G | A | 0.23 | 0.136 | 0.018 | 2.16E-13 |
| glycemic control | rs2876354 | T | C | 0.47 | -0.083 | 0.016 | 3.33E-07 |
| glycemic control | rs7742292 | C | T | 0.4 | 0.041 | 0.016 | 1.04E-02 |
| glycemic control | rs2982521 | T | A | 0.63 | -0.11 | 0.016 | 2.09E-11 |
| glycemic control | rs9390022 | C | T | 0.38 | -0.042 | 0.016 | 8.66E-03 |
| glycemic control | rs1538247 | C | T | 0.3 | 0.093 | 0.017 | 7.76E-08 |
| glycemic control | rs2179168 | A | G | 0.8 | 0.046 | 0.019 | 1.54E-02 |
| glycemic control | rs501470 | G | T | 0.47 | -0.089 | 0.015 | 6.20E-09 |
| glycemic control | rs4709746 | T | C | 0.13 | -0.05 | 0.023 | 2.95E-02 |
| glycemic control | rs7774074 | A | C | 0.21 | 0.039 | 0.019 | 3.98E-02 |
| glycemic control | rs35261542 | A | C | 0.26 | 0.268 | 0.017 | 1.55E-50 |
| glycemic control | rs3117189 | G | A | 0.85 | 0.281 | 0.021 | 3.05E-37 |
| glycemic control | rs7748962 | A | G | 0.77 | 0.113 | 0.018 | 8.41E-10 |
| glycemic control | rs9472139 | C | G | 0.29 | 0.065 | 0.017 | 1.47E-04 |
| glycemic control | rs3798519 | C | A | 0.18 | 0.107 | 0.02 | 1.46E-07 |
| glycemic control | rs9370243 | T | G | 0.08 | 0.079 | 0.028 | 4.82E-03 |
| glycemic control | rs9449295 | C | T | 0.54 | 0.036 | 0.015 | 1.63E-02 |
| glycemic control | rs9379084 | A | G | 0.12 | -0.198 | 0.025 | 1.59E-14 |
| glycemic control | rs187653072 | C | T | 0.03 | 0.134 | 0.044 | 2.38E-03 |
| glycemic control | rs73184014 | G | A | 0.22 | -0.053 | 0.019 | 5.32E-03 |
| glycemic control | rs6976111 | A | C | 0.3 | 0.074 | 0.017 | 1.66E-05 |
| glycemic control | rs13237518 | A | C | 0.41 | 0.048 | 0.016 | 2.75E-03 |
| glycemic control | rs3996350 | C | G | 0.5 | -0.086 | 0.015 | 1.89E-08 |
| glycemic control | rs60251368 | G | A | 0.06 | 0.096 | 0.034 | 4.79E-03 |
| glycemic control | rs4252505 | G | A | 0.06 | 0.07 | 0.031 | 2.38E-02 |
| glycemic control | rs17168486 | T | C | 0.17 | 0.162 | 0.02 | 4.21E-15 |
| glycemic control | rs4725959 | G | A | 0.22 | 0.042 | 0.019 | 2.68E-02 |
| glycemic control | rs10228796 | G | C | 0.55 | 0.16 | 0.015 | 1.33E-24 |
| glycemic control | rs6946660 | C | T | 0.35 | -0.107 | 0.016 | 6.88E-11 |
| glycemic control | rs2188848 | G | A | 0.2 | -0.055 | 0.019 | 3.84E-03 |
| glycemic control | rs860262 | A | C | 0.5 | -0.158 | 0.015 | 4.73E-24 |
| glycemic control | rs917195 | T | C | 0.23 | -0.073 | 0.018 | 5.83E-05 |
| glycemic control | rs730497 | A | G | 0.18 | 0.445 | 0.02 | 4.27E-97 |
| glycemic control | rs73121277 | C | T | 0.28 | 0.084 | 0.017 | 1.12E-06 |
| glycemic control | rs6975279 | A | C | 0.26 | 0.101 | 0.018 | 3.67E-08 |
| glycemic control | rs6956980 | C | T | 0.53 | 0.083 | 0.015 | 5.56E-08 |
| glycemic control | rs7834323 | C | T | 0.29 | -0.074 | 0.017 | 1.66E-05 |
| glycemic control | rs727582 | G | A | 0.34 | -0.093 | 0.016 | 1.22E-08 |
| glycemic control | rs13266634 | T | C | 0.31 | -0.277 | 0.017 | 9.11E-54 |
| glycemic control | rs12056338 | T | G | 0.42 | 0.05 | 0.016 | 1.83E-03 |
| glycemic control | rs1561927 | T | C | 0.73 | -0.048 | 0.017 | 4.79E-03 |
| glycemic control | rs35753840 | C | A | 0.39 | 0.054 | 0.016 | 7.78E-04 |
| glycemic control | rs13268508 | T | C | 0.38 | 0.087 | 0.016 | 9.23E-08 |
| glycemic control | rs2953845 | T | C | 0.55 | 0.042 | 0.015 | 5.15E-03 |
| glycemic control | rs6558173 | T | G | 0.35 | 0.039 | 0.016 | 1.47E-02 |
| glycemic control | rs2725370 | C | T | 0.7 | -0.049 | 0.017 | 3.99E-03 |
| glycemic control | rs57735787 | G | A | 0.25 | -0.042 | 0.018 | 1.95E-02 |
| glycemic control | rs13262861 | A | C | 0.17 | -0.121 | 0.021 | 1.61E-08 |
| glycemic control | rs7813865 | C | T | 0.29 | 0.041 | 0.017 | 1.58E-02 |
| glycemic control | rs10101067 | C | G | 0.08 | 0.092 | 0.029 | 1.56E-03 |
| glycemic control | rs28792187 | G | A | 0.07 | 0.123 | 0.03 | 4.86E-05 |
| glycemic control | rs1895874 | A | G | 0.5 | 0.048 | 0.015 | 1.42E-03 |
| glycemic control | rs10808671 | G | A | 0.53 | -0.073 | 0.015 | 1.61E-06 |
| glycemic control | rs60384372 | G | A | 0.47 | -0.056 | 0.015 | 2.09E-04 |
| glycemic control | rs1567353 | G | C | 0.31 | 0.035 | 0.017 | 3.92E-02 |
| glycemic control | rs10119430 | A | G | 0.79 | -0.054 | 0.019 | 4.53E-03 |
| glycemic control | rs1431819 | G | A | 0.7 | 0.038 | 0.017 | 2.52E-02 |
| glycemic control | rs10818763 | T | C | 0.13 | -0.108 | 0.023 | 3.58E-06 |
| glycemic control | rs10739629 | T | C | 0.51 | -0.036 | 0.015 | 1.63E-02 |
| glycemic control | rs529565 | C | T | 0.32 | 0.164 | 0.017 | 1.52E-20 |
| glycemic control | rs28642213 | G | A | 0.75 | 0.169 | 0.018 | 1.41E-19 |
| glycemic control | rs12380322 | G | A | 0.39 | 0.051 | 0.016 | 1.49E-03 |
| glycemic control | rs10965247 | G | A | 0.18 | -0.302 | 0.02 | 1.27E-46 |
| glycemic control | rs7018475 | G | T | 0.26 | 0.178 | 0.018 | 1.79E-21 |
| glycemic control | rs11788619 | T | A | 0.03 | -0.134 | 0.048 | 5.28E-03 |
| glycemic control | rs2150854 | T | G | 0.33 | 0.072 | 0.016 | 8.71E-06 |
| glycemic control | rs4237150 | C | G | 0.4 | 0.091 | 0.016 | 2.43E-08 |
| glycemic control | rs67269808 | G | A | 0.06 | -0.13 | 0.032 | 5.67E-05 |
| glycemic control | rs2796441 | A | G | 0.42 | -0.096 | 0.016 | 4.24E-09 |
| glycemic control | rs7023781 | T | C | 0.27 | 0.058 | 0.017 | 6.83E-04 |
| glycemic control | rs10993072 | T | C | 0.32 | 0.083 | 0.016 | 3.33E-07 |
| glycemic control | rs28496034 | G | C | 0.33 | -0.057 | 0.016 | 3.96E-04 |

**eTable 4. the main results of significant risk factors in univariable MR.**

| **outcome** | **exposure** | **method** | **nsnp** | **pval** | **or** | **or_lci95** | **or_uci95** |
| --- | --- | --- | --- | --- | --- | --- | --- |
| LS | PP | Simple median | 76 | 0.032 | 1.05 | 1.00 | 1.10 |
| LS | PP | Weighted median | 76 | 0.338 | 1.02 | 0.97 | 1.08 |
| LS | PP | MR Egger | 76 | 0.888 | 1.01 | 0.89 | 1.14 |
| LS | PP | IVW (fixed effects) | 76 | 0.012 | 1.04 | 1.01 | 1.08 |
| LS | DBP | Simple median | 75 | 0.023 | 1.07 | 1.01 | 1.14 |
| LS | DBP | Weighted median | 75 | 0.021 | 1.07 | 1.01 | 1.14 |
| LS | DBP | MR Egger | 75 | 0.606 | 0.96 | 0.82 | 1.12 |
| LS | DBP | IVW (fixed effects) | 75 | 0.003 | 1.06 | 1.02 | 1.11 |
| LS | SBP | Simple median | 98 | 0.010 | 1.04 | 1.01 | 1.08 |
| LS | SBP | Weighted median | 98 | 0.002 | 1.05 | 1.02 | 1.08 |
| LS | SBP | MR Egger | 98 | 0.036 | 1.12 | 1.01 | 1.24 |
| LS | SBP | IVW (fixed effects) | 98 | 4.64E-07 | 1.06 | 1.03 | 1.08 |
| LS | HDL cholesterol \|\| id:ieu-b-109 | Simple median | 257 | 0.144 | 0.87 | 0.72 | 1.05 |
| LS | HDL cholesterol \|\| id:ieu-b-109 | Weighted median | 257 | 0.390 | 0.92 | 0.77 | 1.11 |
| LS | HDL cholesterol \|\| id:ieu-b-109 | MR Egger | 257 | 0.111 | 0.87 | 0.73 | 1.03 |
| LS | HDL cholesterol \|\| id:ieu-b-109 | IVW (multiplicative random effects) | 257 | 0.008 | 0.86 | 0.76 | 0.96 |
| LS | apolipoprotein A-I \|\| id:ieu-b-107 | Simple median | 226 | 0.217 | 0.89 | 0.73 | 1.07 |
| LS | apolipoprotein A-I \|\| id:ieu-b-107 | Weighted median | 226 | 0.554 | 0.95 | 0.79 | 1.13 |
| LS | apolipoprotein A-I \|\| id:ieu-b-107 | MR Egger | 226 | 0.352 | 0.91 | 0.76 | 1.10 |
| LS | apolipoprotein A-I \|\| id:ieu-b-107 | IVW (multiplicative random effects) | 226 | 0.034 | 0.88 | 0.78 | 0.99 |
| LS | triglycerides \|\| id:ieu-b-111 | Simple median | 234 | 0.001 | 1.45 | 1.18 | 1.80 |
| LS | triglycerides \|\| id:ieu-b-111 | Weighted median | 234 | 0.386 | 1.08 | 0.91 | 1.29 |
| LS | triglycerides \|\| id:ieu-b-111 | MR Egger | 234 | 0.861 | 1.02 | 0.85 | 1.21 |
| LS | triglycerides \|\| id:ieu-b-111 | IVW (multiplicative random effects) | 234 | 0.028 | 1.14 | 1.01 | 1.29 |
| LS | apolipoprotein B \|\| id:ieu-b-108 | Simple median | 142 | 0.190 | 1.14 | 0.94 | 1.39 |
| LS | apolipoprotein B \|\| id:ieu-b-108 | Weighted median | 142 | 0.546 | 1.06 | 0.88 | 1.26 |
| LS | apolipoprotein B \|\| id:ieu-b-108 | MR Egger | 142 | 0.377 | 1.09 | 0.90 | 1.31 |
| LS | apolipoprotein B \|\| id:ieu-b-108 | IVW (multiplicative random effects) | 142 | 0.030 | 1.15 | 1.01 | 1.31 |
| LS | Type 2 diabetes | Simple median | 113 | 0.001 | 1.16 | 1.06 | 1.26 |
| LS | Type 2 diabetes | Weighted median | 113 | 0.007 | 1.15 | 1.04 | 1.28 |
| LS | Type 2 diabetes | MR Egger | 113 | 0.190 | 1.09 | 0.96 | 1.24 |
| LS | Type 2 diabetes | IVW (fixed effects) | 113 | 1.02E-04 | 1.12 | 1.06 | 1.18 |
| LS | Height | Simple median | 290 | 0.204 | 0.92 | 0.81 | 1.05 |
| LS | Height | Weighted median | 290 | 0.278 | 0.93 | 0.81 | 1.06 |
| LS | Height | MR Egger | 290 | 0.034 | 0.76 | 0.59 | 0.98 |
| LS | Height | IVW (multiplicative random effects) | 290 | 0.011 | 0.88 | 0.81 | 0.97 |
| LS | Education | Simple median | 30 | 0.075 | 0.59 | 0.33 | 1.05 |
| LS | Education | Weighted median | 30 | 0.084 | 0.58 | 0.32 | 1.07 |
| LS | Education | MR Egger | 30 | 0.568 | 0.46 | 0.03 | 6.56 |
| LS | Education | IVW (fixed effects) | 30 | 0.006 | 0.55 | 0.36 | 0.84 |
| LS | Fasting proinsulin | Simple median | 9 | 0.030 | 2.19 | 1.08 | 4.45 |
| LS | Fasting proinsulin | Weighted median | 9 | 0.024 | 1.80 | 1.08 | 3.00 |
| LS | Fasting proinsulin | MR Egger | 9 | 0.517 | 1.29 | 0.62 | 2.66 |
| LS | Fasting proinsulin | IVW (fixed effects) | 9 | 0.012 | 1.54 | 1.10 | 2.15 |
| LS | Fibrinogen | Simple median | 33 | 0.245 | 2.13 | 0.59 | 7.65 |
| LS | Fibrinogen | Weighted median | 33 | 0.153 | 2.36 | 0.73 | 7.67 |
| LS | Fibrinogen | MR Egger | 33 | 0.171 | 5.13 | 0.52 | 50.58 |
| LS | Fibrinogen | IVW (fixed effects) | 33 | 0.020 | 2.52 | 1.16 | 5.47 |
| LS | Atrial fibrillation | Simple median | 74 | 0.357 | 0.95 | 0.85 | 1.06 |
| LS | Atrial fibrillation | Weighted median | 74 | 0.288 | 0.95 | 0.85 | 1.05 |
| LS | Atrial fibrillation | MR Egger | 74 | 0.111 | 0.90 | 0.79 | 1.02 |
| LS | Atrial fibrillation | IVW (fixed effects) | 74 | 0.010 | 0.92 | 0.86 | 0.98 |

| **eTable 5. Multivariable MR estimates for blood pressure and lipids.** | | | | | |  |
| --- | --- | --- | --- | --- | --- | --- |
| **Outcome** | **Method** | **SNP N** | **Exposure** | **BETA** | **SE** | **p-value** |
| ls | IVW | 255 | DBP | 0.082 | 0.235 | 0.729 |
|  |  |  | SBP | -0.024 | 0.239 | 0.919 |
|  |  |  | PP | 0.064 | 0.241 | 0.789 |
|  | MR-Egger | 255 | DBP | 0.08 | 0.237 | 0.735 |
|  |  |  | SBP | -0.025 | 0.239 | 0.918 |
|  |  |  | PP | 0.065 | 0.241 | 0.788 |
| ls | IVW | 384 | ApoB | 0.171 | 0.296 | 0.564 |
|  |  |  | LDL | -0.136 | 0.315 | 0.665 |
|  |  |  | TG | 0.196 | 0.071 | 0.005* |
|  | MR-Egger | 384 | ApoB | 0.14 | 0.295 | 0.636 |
|  |  |  | LDL | -0.192 | 0.316 | 0.543 |
|  |  |  | TG | 0.151 | 0.075 | 0.045* |
|  | MR-Lasso | 357 | ApoB | -0.006 | 0.261 | 0.98 |
|  |  |  | LDL | 0.035 | 0.28 | 0.902 |
|  |  |  | TG | 0.261 | 0.063 | 0.000* |
| ls | IVW | 435 | ApoAI | 0.023 | 0.187 | 0.900 |
|  |  |  | HDL | -0.201 | 0.178 | 0.26 |
|  | MR-Egger | 435 | ApoAI | 0.065 | 0.194 | 0.738 |
|  |  |  | HDL | -0.187 | 0.179 | 0.296 |
|  | MR-Lasso | 408 | ApoAI | -0.046 | 0.167 | 0.783 |
|  |  |  | HDL | -0.149 | 0.161 | 0.354 |

| **eTable 6. MR-Egger intercept and Q test result in the study.** | | | | | | |
| --- | --- | --- | --- | --- | --- | --- |
| **outcome** | **exposure** | **IVW-Q test** | | **Egger regression (intercept)** | | |
|  |  | **Q-statistic** | **Q_pval** | **egger_intercept** | **se** | **pval** |
| **univariable MR** |  |  |  |  |  |  |
| lacunar stroke | PP | 90.6 | 0.106 | 0.005 | 0.010 | 0.589 |
| lacunar stroke | DBP | 90.0 | 0.099 | 0.014 | 0.011 | 0.178 |
| lacunar stroke | SBP | 106.6 | 0.238 | -0.013 | 0.012 | 0.256 |
| lacunar stroke | HDL | 326.9 | 0.002 | 0.000 | 0.003 | 0.873 |
| lacunar stroke | apolipoprotein A-I | 289.5 | 0.002 | -0.001 | 0.003 | 0.614 |
| lacunar stroke | triglycerides | 313.7 | 0.000 | 0.005 | 0.003 | 0.073 |
| lacunar stroke | apolipoprotein B | 195.6 | 0.002 | 0.003 | 0.004 | 0.394 |
| lacunar stroke | Type 2 diabetes | 291.8 | 0.142 | 0.004 | 0.003 | 0.153 |
| lacunar stroke | Height | 391.2 | 0.000 | 0.005 | 0.004 | 0.209 |
| lacunar stroke | Education | 20.0 | 0.894 | 0.003 | 0.024 | 0.891 |
| lacunar stroke | Fasting proinsulin | 11.2 | 0.190 | 0.009 | 0.015 | 0.578 |
| lacunar stroke | Fibrinogen | 63.6 | 0.001 | -0.008 | 0.011 | 0.491 |
| lacunar stroke | Atrial fibrillation | 108.0 | 0.083 | 0.004 | 0.005 | 0.405 |
| **multivariable MR** |  |  |  |  |  |  |
| lacunar stroke | blood pressure: PP SBP PP | 287.6 | 0.061 | 0.000 | 0.003 | 0.951 |
| lacunar stroke | lipids: ApoB LDL TG | 524.0 | 0.000 | 0.003 | 0.002 | 0.085 |
| lacunar stroke | lipids: apolipoprotein A-I HDL | 550.2 | 1.00E-04 | -0.002 | 0.002 | 0.434 |
